# Towards Clinical-Grade Bioengineered Airways: A Study on Stem Cell Renewal and Epithelial Differentiation

**DOI:** 10.1101/2023.12.28.573530

**Authors:** Adamo Davide, Genna Vincenzo Giuseppe, Galaverni Giulia, Chiavelli Chiara, Merra Alessia, Lepore Fabio, Boraldi Federica, Quaglino Daniela, Evangelista Jessica, Lococo Filippo, Pellegrini Graziella

## Abstract

**Background:** Despite their life-saving potential, tissue engineering approaches for the treatment of extensive tracheal and bronchial defects still face significant limitations. A major challenge is the inability to regenerate a functional airway epithelium containing the appropriate amount of stem cells required for long-term tissue renewal following transplantation of the bioengineered graft. In this scenario, validation assays and quality controls are needed to guide each step of the regeneration process.

**Methods:** Stem cell depletion is often due to suboptimal culture conditions, therefore we tested the ability of a clinical-grade culture system to support the safe and efficient in vitro expansion and differentiation of primary human tracheal and bronchial epithelial cells. Single-cell clonal analysis was used to unravel the heterogeneity of airway basal cells and to understand tissue-specific regeneration and differentiation mechanisms. Functional assays were used to investigate the wound healing ability and tightness of the regenerated epithelium under the selected culture conditions.

**Results:** Primary tracheobronchial epithelial cells showed an impressive proliferative potential, allowing the regeneration of a mature and functional epithelium without immortalisation events. Analysis at the single cell level allowed the identification of the subpopulation of basal cells endowed with *in vitro* self-renewal, distinguishing them from transient amplifying cells. This approach has further defined the hierarchy of cellular differentiation and its correlation with regenerative and differentiation potential.

**Conclusions:** Our results show that primary airway epithelial cell cultures can maintain stem cells together with their differentiation lineages *in vitro*. Airway cells can be safely and effectively used in autologous tissue engineering approaches when cultured under appropriate and well-standardised conditions. In addition to the validation assays proposed for the development of new advanced therapy products, this study outlines possible quality controls to enhance therapeutic success and maximise patient safety in future clinical applications.

## Background

Knowledge of the importance of the respiratory tract dates back to 3,600 BC, with tracheotomy operations depicted in Egyptian tablets, as the respiratory system is one of the most important and intricate systems in the body [1,2]. This field has garnered interest owing to several pathological conditions, such as malignancies, congenital malformations, infections, inflammation, or postoperative complications, that can profoundly alter airway functions, endangering the patient’s life [3–7]. Over the years, extensive circumferential tracheal or bronchial damages have posed a significant challenge to airway medicine [8,9]. In these cases, conventional surgical approaches have proven ineffective, highlighting the pressing demand for innovative reconstructive strategies [10–13].

Tissue engineering (TE) has been investigated as a promising solution for replacing damaged or pathological airway tissue [5,14]. However, the clinical outcomes of TE applications have highlighted their limitations, including the need for biocompatible supporting materials with appropriate biomechanical properties [11,13,15,16], immune rejection risks [17], delayed vascularization of the bioengineered graft [18] and, above all, the inability to regenerate a functional and self-renewing airway epithelium on the engineered implant in the long term [8,19–21].

The human airway is a remarkable structure formed by the coordinated development of endodermal and mesodermal tissues [22]. Impairment of airway epithelial function leads to complications such as mucus build-up, infection, graft stenosis, granulation tissue formation, chondromalacia and fibrogenic reactions [8,20,21,23], with profound physical and emotional challenges for patients.

The airway epithelium relies on various primary cell populations, including basal, goblet, club, and ciliated cells, and other less abundant cell types, collectively accounting for 0.8% of the total airway epithelium composition [24]. The efficient expansion of primary cells from airway epithelia, maintenance of tissue architecture, and inherent heterogeneity to the extent required for transplantation are major challenges for airway TE [25–27].

Indeed, preclinical studies describing primary airway epithelial cell cultivation *in vitro* have generally reported a slow cell growth rate [20] and minimal proliferative potential, with a very limited number of subculturing passages [28]. Beyond this limit, cells lose their capacity to proliferate, produce mucus, and express tight junction proteins [29–31], resulting in unbalanced differentiation. Consequently, autologous airway epithelial cells have been reported to be inadequate for clinical regeneration and unreliable for TE approaches [20]. Maintaining the epithelial stem cell population is of utmost importance to promptly achieve effective *ex vivo* expansion and differentiation, which is required for life-saving reconstructions [32]. Stem cell preservation must be ensured in all stages of the engineering process, from biopsy collection and cell extraction to *in vitro* expansion and graft repopulation, and finally through a well-defined vascularization strategy that ensures sufficient blood supply to the transplanted tissue for long-term therapeutic ends [8,10,31].

Based on successful clinical applications of several regenerative medicine approaches, permanent engraftment and long-term regeneration of transplanted epithelial tissues appear to be dependent on an adequate number of stem cells [33–36]. Hence, identifying specific biological parameters that can unequivocally determine the quality of epithelial culture [37–40] is critical and required by regulatory authorities to approve TE clinical applications.

Here, we demonstrated the selection of a clinical-grade culture system [33,41–43] able to achieve efficient, rapid and safe expansion of human primary airway tracheal (AT) and bronchial (AB) epithelial cells while preserving their tissue-specific heterogeneity. The main goals of this study were i) to maintain the long-term proliferative potential of airway epithelial cells and their multipotency by expressing the major differentiation lineages, ii) to identify and characterize airway epithelial stem cells, which are essential for self-maintenance of the tissue *in vivo*, and iii) to provide a basis for quality controls in bioengineering processes. The prospective identification of specific biomarkers can be relevant for diagnosis and follow-up *in vivo.* Additionally, AT and AB epithelial cell differentiation kinetics unravelled the different roles of stem cells and transient amplifying progenitors in maintaining airway epithelial heterogeneity and potency. Increased understanding of tissue-specific biology and regenerative potential under *in vitro* expansion will help overcome hurdles in airway TE and stem cell maintenance, promoting successful translational research.

## Methods

### Human samples

Human tracheal and bronchial samples were collected from n=5 male donors and n=5 female donors aged between 38 and 76 years after informed consent and ethical committee approval. Exclusion criteria implied patients suffering from neoplastic diseases or infections such as aspergillosis, nocardiosis, tuberculosis or SARS-CoV-2. Human skin samples were obtained from healthy living donors as a byproduct of surgical waste (abdominoplasty or mammoplasty) upon informed consent and compliance with Italian regulations.

### Cell lines and primary cell culture

After collection in the surgery room, tracheal and bronchial biopsies were shipped from the clinical centres to the “Centre for Regenerative Medicine of Modena and Reggio-Emilia, Modena, Italy” through a clinically validated, temperature-controlled transport packaging. After transport within a clinical-grade (CG) culture medium and no more than 36 hours after sample collection, the biopsies were minced and dissociated with 0.05% trypsin (Thermo Fisher Scientific, Waltham, Massachusetts, USA) and 0.01% 2,2′,2″,2‴-(Ethane-1,2-diyldinitrilo) tetra-acetic acid (EDTA) at 37°C for 2‒6 hours. Every 30 min, epithelial cells were collected, plated (2.5–3 × 10^4^/cm^2^) on Good Manufacturing Practice (GMP)-certified lethally irradiated 3T3-J2 cells (2.4 × 10^4^/cm^2^), and grown at 37°C, 5% CO_2_ in 95% humidified atmosphere in Dulbecco’s modified Eagle’s (DMEM) and Ham’s F12 media (2:1 mixture, (EuroClone, Milan, Italy), 10% fetal bovine serum (FBS, Thermo Fisher Scientific), 50 lU/ml penicillin–streptomycin (EuroClone, Milan, Italy), 4 mM glutamine (EuroClone, Milan, Italy), 0.18 mM adenine (Merck, Darmstadt, Germany), 5 mg/ml insulin (Eli Lilly and Company, Indianapolis, Indiana, USA), 0.1 nM cholera toxin (List Biological, California, USA), 0.4 mg/ml hydrocortisone (Avantor, Pennsylvania, USA), 2 nM triiodothyronine (Peptido Gmbh). At the first feeding time, three days after plating, 10 ng/ml epidermal growth factor (EGF, Austral Biologicals, California, USA) was added to the medium. Primary cultures were fed every other day. When airway cultures reached subconfluence, the epithelial cells were dissociated at 37°C for 15 min in trypsin and either frozen at -196°C or serially propagated at a density of 6x10^6^ cells/cm^2^ until replicative senescence. All the reagents used for cell extraction and culture were extensively selected and screened for contaminants and the absence of cytotoxic effects.

Mouse 3T3-J2 cells, used as feeder layer (FL) for human epithelial cells, were cultivated in DMEM supplemented with 10% gamma-irradiated donor adult bovine serum, penicillin–streptomycin (50 IU/ml) and glutamine (4 mM) and detached with 0.25% Trypsin-EDTA 1X (Thermo Fisher Scientific, Waltham, Massachusetts, USA). Mouse 3T3-J2 cells were originally received as a gift from Prof. Howard Green, Harvard Medical School (Boston, MA, USA), and a GMP-certified master cell bank was established.

### Colony-forming efficiency assay, population doublings and growth rate (doubling time)

The colony-forming efficiency (CFE) was calculated after cell extraction from human biopsies and at each serial passage of tracheobronchial lifespans from the indicator dish stained with rhodamine B (Sigma‒Aldrich, St. Louis, Missouri, USA) after 12 days of cultivation. Briefly, a small aliquot of cells - ranging from 200 to 2000 - was plated onto indicator culture dishes previously seeded with 3T3-J2 FL and cultured following the abovementioned protocol. Under examination with a dissecting microscope, colonies were scored as progressively growing or aborted, as previously described [44,45]. The percentage of clonogenic cells corresponds to the number of grown colonies on the total number of plated cells; the percentage of aborted colonies was calculated as the number of aborted colonies on the total number of grown colonies. The following formula was used to score the number of cell doublings x = 3.322 log N/No, where N is the total number of epithelial cells collected at each passage and No is the number of clonogenic cells plated. Doubling time was calculated using the following formula: number of hours to reach subconfluence/number of cell doublings performed, considering the passages with standard cell number seeding and above 2 .5% clonogenicity.

### Clonal analysis

For the clonal analysis, subconfluent airway cultures were trypsinized and, after limiting dilution, single cells were inoculated into 96-well plates (1 cell per well) coated with 3T3-J2 FL cells. After seven days of culture, wells containing single clones were selected under an inverted microscope , and each clone was photographed and then dissociated through trypsin digestion to obtain a single-cell suspension. Afterwards, each cell suspension was divided into three or four parts. Typically, one-quarter of the clone was plated into an indicator dish for CFE analysis, whereas the other three quarters were used in various experiments to characterize each clone’s regenerative potential. CFE assay allows clonal classification according to the number of aborted colonies. Briefly, the clones that generated between 0-6% of aborted colonies were classified as holoclones (H), the clones that generated between 6-95% of aborted colonies were classified as meroclones (M), and the clones that generated more than 95% of aborted colonies were classified as paraclones (P). Among meroclones, a further classification distinguished them in early meroclones (EM, 6-20% of aborted colonies), intermediate meroclones (IM, 20-60% of aborted colonies) and late meroclones (LM, 60-95% of aborted colonies). The clone size was calculated via a ×10 objective and ZEN plug-in AxioVision SE64 software (Rel. 4.9.1, Zeiss, Oberkochen, Germany).

### Unclustered analysis of airway clonogenic cells

Subconfluent airway epithelial cultures were enzymatically detached and, after limiting dilution, single cells were inoculated into 96-well plates containing 3T3-J2 FL cells. On the 7th day of culture, single AT and AB clones were fixed with 3% paraformaldehyde (10 min at RT) or cold methanol (10 min at -20°C) and immunostained as described in the “Immunofluorescence staining” section. Images were acquired through Cell Observer Z.1 microscope (Zeiss) and ZEN plug-in AxioVision software (Rel. 4.8, Zeiss, Oberkochen, Germany).

### *In vitro* self-renewal assay

To evaluate the self-renewal potential of the different clonal categories under stressful conditions, AT and AB clones grown as described in the “clonal analysis” section were detached and concomitantly seeded into an indicator dish for clonal type classification (one-third of the clone) and in a well of a 24-well plate for 28/30 days coated with FL (two-thirds of the clone). During this prolonged culture period, the CG cell culture medium was first changed on day three and then changed daily until the 28/30^th^ day of culture. The progeny of each clone were subsequently detached, and CFE assay was performed. Specifically, we plated 5,000–10,000 cells for H (to allow a precise colony count), whereas for the other clonal types (M and P), we plated all the detached cells in indicator dishes to provide an overall assessment of residual clonogenicity.

### Air-liquid interface (ALI) culture

For the ALI culture, tracheal and bronchial epithelial cells derived from subconfluent cultures were seeded (2.4 × 10^4^/cm^2^) onto a human de-epithelialized derma (used as a biological scaffold) coated with lethally irradiated 3T3‒J2 FL cells and located within Millicell^®^ culture plate inserts (Merck Millipore, Burlington, MA, USA). Human AT and AB epithelial cells were cultured onto this matrix submerged in CG culture medium for seven days, after which the medium was removed from the apical chamber to start the air-liquidALI culture. The culture medium was changed every other day until day 28-30. At the end of this period, airlifted cultures were embedded in optimal cutting temperature (OCT, Bio-Optica) compound and stored at -80°C until use. The same protocol was applied for the ALI culture of the clonal progenies, with the only exception being the seeding density. Indeed, for that experiment, three-quarters of each clone was used to establish ALI cultures.

### De-epithelialization of the human dermis

To de-epithelialize the human dermis, human skin biopsies derived from abdominoplasty or mammoplasty were subjected to thermal shock. Specifically, the samples were submerged in preheated 1X PBS at 60°C for 30 seconds and then immediately transferred to a container with 1X PBS at 4°C for one minute. Following this treatment, the epidermis was carefully separated from the dermis via sterile forceps, with care taken to preserve the integrity of the dermal tissue.

### Immunofluorescence staining

For immunofluorescence analysis, tracheal and bronchial human specimens and the ALI cultures on human de-epithelialized dermis were embedded in OCT compound (Bio-Optica) and stored at -80°C. Subsequently, 8-20 µm-thick sections were cut with a cryostat and mounted onto glass slides. These sections, as well as airway epithelial cultures grown onto glass slide coverslips (15,000 cells/well of a 24-well plate) and single clones grown in 96-well plates, were fixed with 3% paraformaldehyde (10 min at RT) or cold methanol (10 min at -20°C). The samples were then carefully washed with 1× PBS, permeabilized with 0.5–1% Triton X-100 in 1× PBS (10 min at RT) and subsequently blocked with blocking solution (2% BSA in PBS) for 30 min at 37°C or 60 min at RT. The samples were first incubated for 30 min at 37°C with the primary antibody and later with the appropriate secondary antibody (typically Alexa Fluor conjugates, Thermo Fisher Scientific, **Supplementary Table 1**) for 30 min at 37°C, both of which were diluted in blocking solution. The following primary antibodies were used: rabbit polyclonal anti-KERATIN 5 (1:1000, Biolegend), rabbit polyclonal anti-KERATIN 14 (1:1000, Biolegend), mouse monoclonal anti-KERATIN 18 (1:100, Abcam), rabbit monoclonal anti-KERATIN 19 (1:2000, Abcam), mouse monoclonal anti-Involucrin (1:100, Leica Microsystems), mouse monoclonal anti-Acetylated Tubulin (1:100, Sigma), mouse monoclonal anti-mucin (1:20, Progen), rat monoclonal anti-Uteroglobin (1:20, R&D System), rabbit polyclonal anti-ZO-1 (1:100, Thermo Fisher Scientific), rabbit polyclonal anti-Ki-67 (1:100, Abcam), rabbit polyclonal anti-p63-alpha (1:5000, custom), and mouse monoclonal anti-14-3-3- Sigma (1:200, Abcam). The cell nuclei were stained with 4’,6-diamidino-2-phenylindole (DAPI) for 5 min at RT, and the coverslips were mounted with fluorescent mounting medium (Dako, Agilent Technologies, Santa Clara, CA, USA). Images of *the in vivo* sections, airlifted culture sections, and airway epithelial cultures grown onto glass slide coverslips were acquired with a laser-scanning confocal microscope (LSM 900, Zeiss, Oberkochen, Germany) and ZEN 2 software (Rel. 3.3, Zeiss, Oberkochen, Germany). A Zeiss EC PlanNeofluar 40x/1.3 oil immersion objective was used to visualize the fluorescent signals. Instead, images of single clones grown in 96-well plates were acquired via a Z1 motorized inverted fluorescence microscope (Zeiss Axio Observer Z1, Zeiss, Oberkochen, Germany). Per each single clone, the staining for the analysed marker was annotated as positive or negative. Then, the ratio between the number of positive clones and the total number of analysed clones was calculated.

### Airway specialized cell abundance in lifespan serial passages

To determine the abundance of goblet and club cells within serial passages of AT and AB cultures, one entire coverslip from each analysed passage of the cultures was stained for MUC5AC and Uteroglobin (**Supplementary Table 1**) and thoroughly screened (from top to bottom) for the presence of these cells. Owing to varying expression levels of the aforementioned proteins and the natural stratification of the epithelial colonies, precise quantification of the two cell populations was unattainable. Instead, we assigned a score to each analysed culture based on the abundance of positive cells. The grading criteria were as follows: -, no positive cells were present (none); +, fewer than 10 positive cells present (low); ++, positive cells present in approximately 30% of the colonies (moderate); +++, positive cells present in approximately 60% of the colonies (high); and ++++, positive cells present in approximately 90% of the colonies (very high). For each lifespan interval (I-V), multiple cultures on glass slide coverslips were analysed. Therefore, the grading for each interval was represented as the mean value of these analyses.

### Western blotting

For the WB analysis conducted on the serial passages of AT and AB culture lifespans, proteins were extracted from dry pellets via RIPA buffer (Sigma-Aldrich, St. Louis, Missouri, USA; cat# R0278) supplemented with phosphatase and protease inhibitor cocktails (Thermo Fisher Scientific, Waltham, MA, USA; cat# 78420; cat# 72426) at 0–4°C. The Bradford assay (Bio-Rad, Hercules, CA, USA; cat# 500-0006) and the spectrophotometer were used to quantify the total protein amount. Equal amounts of protein were electrophoresed on NuPage 4-12% Bis-Tris protein gels (Thermo Fisher Scientific, St. Louis, Missouri, USA; cat# NP0322) and transferred to nitrocellulose membranes (Merck-Millipore, Burlington, MA, USA; cat#1620115) at 100 V and 4°C for 2 hours. The membranes were treated with blocking solution containing 5% (w/v) nonfat milk (Bio-Rad, Hercules, CA, USA, cat#1706404) in 0.01% (v/v) Tween-20 (Sigma-Aldrich, St. Louis, Missouri, USA, cat#P1379) in 1× PBS. Protein band immunoreactions were performed with different primary antibodies (**Supplementary Table 1**) diluted in blocking solution, which were added to the membranes overnight at 4°C.

The following primary antibodies were used: rabbit polyclonal anti-p63-alpha (1:5000, customized), rabbit monoclonal anti-BMI1 (1:500, Cell Signaling Technology), rabbit monoclonal anti-SOX2 (1:200, Cell Signaling Technology), mouse monoclonal anti-GAPDH (1:10.000, Abcam), mouse monoclonal anti-vinculin (1:10.000, Sigma-Aldrich), rabbit polyclonal anti-KERATIN 14 (1:40.000, Biolegend), mouse monoclonal anti-Involucrin (1:10.000, Leica Microsystems), and mouse monoclonal anti-actin (1:5000, Abcam).

The corresponding HRP-conjugated secondary antibodies (**Supplementary Table 1**) were diluted in blocking solution and incubated for 1 hour at RT after three washes with 1× PBS. Protein signals were developed via a chemiluminescent labelling reagent (Supersignal West Pico Chemiluminescent Substrate, Thermo Scientific, cat#34580) and acquired with ChemiDocTM (Bio-Rad, Hercules, CA, USA) and Image Lab software (Bio-Rad, Hercules, CA, USA), while bands quantification was performed via ImageJ software. A grey background on the images was homogeneously added for graphical purposes. For the WB analysis of AT and AB lifespan, each lifespan was subdivided into five consecutive temporal intervals (I-V) to compare markers’ expression levels of the different strains (AT n=3 and AB n =3) in the same airway district. In the case of multiple pellets analysed within the same interval, the mean expression value was considered. Finally, the average and SD of n = 3 AT and n = 3 AB were plotted per time interval, and the expression level of each independent strain was indicated with different shapes. For the WB analysis of clonal progeny, the expression values of H, EM, IM, LM and P of both AT and AB clones are shown for each strain as the mean of the different clones analysed, indicated with dots.

### Scanning electron microscopy (SEM)

AT and AB airlifted cultures were fixed in 2.5% (v/v) glutaraldehyde in 0. 1 M sodium cacodylate for 1 hour at 4°C, washed with 0.1 M sodium cacodylate and postfixed in 1% (w/v) OsO4 for 1 hour at room temperature. The samples were dehydrated in an ascending series of ethanol and critical point dried through liquid carbon dioxide. The specimens were mounted on aluminium stubs and coated with a gold sputter. Finally, the samples were observed under a NOVA NanoSEM 450, and images were acquired in high-vacuum mode with a TLD detector operated at 8 kV.

### Wound healing assay

To perform the wound healing assay, human AT and AB epithelial cells were plated into rectangular cell culture plates (from Cell Comb Scratch Assay, Merck-Millipore, Burlington, MA, USA) and cultivated up to confluence either with CG culture medium and 3T3-J2 FL or with the commercially available defined medium Bronchial Epithelial Cell Growth Medium BulletKit™ (BEGM™, Lonza, Basel, Switzerland). After two days of confluence, the CG culture medium and the BEGM™ were removed, and Cell Combs (from Cell Comb Scratch Assay, Merck-Millipore, Burlington, MA, USA) were used to create two perpendicular scratches across the monolayer. The scratched epithelia were then rinsed with 1× PBS, submerged in CG or BEGM™ culture medium and placed inside the incubator of Cell Observer Z.1 microscope (Zeiss, Oberkochen, Germany) to measure progressive wound healing. Images were taken every 10 min for 14 hours and then every 300 min up to 40 hours. Progressive wound healing was assessed by calculating the area of wound closure through AxioVision Software (version 4.8).

### Trans-epithelial electrical resistance measurement

Transepithelial electrical resistance (TEER) was measured for human tracheal and bronchial epithelia cultured on Millicell^®^ inserts in ALI conditions on days 3, 9, 15, 23, and 29. TEER readings were obtained via a Millicell^®^ ERS-2 voltmeter (Electrical Resistance System, Merck-Millipore, Burlington, MA, USA). The apical surface of cultured epithelia was submerged with CG culture medium only for the time needed to perform the measurement. The resistance values of the replicates were normalized by subtracting the corresponding value of TEER measured in control inserts that did not contain cells. The data were calculated as mean ± SD of the measurements performed at each time point.

### Quantification and statistical analysis

The results are presented as mean ± SD. The significance of differences was analysed by Student’s unpaired two-sample t test via Prism 8 software (version 8.4.0, GraphPad Software, San Diego, CA, USA), and p<0.05 was considered significant. Details about the statistical analyses for each experiment are outlined in the figure legends.

## Results

### Clinical grade culture system

The efficient culture system developed by Rheinwald and Green [46] for *ex vivo* expansion and clinical application of cultured epidermis did not meet current regulatory requirements and was subsequently modified and used for the regeneration of several epithelial tissues [34,42,43,47]. This method has never been studied in human airway epithelial cells or efficiently adapted and standardized for this tissue. Here, we selected conditions in accordance with EU Advanced Therapy Medicinal Products (ATMP) regulations and determined their effects on the airway epithelium (see **Table 1** and Methods).

**Table 1.** Specificities of the clinically-validated culture system and compliance with EU regulations.

| Peculiarities | Specificities | Implications |
| --- | --- | --- |
| Safety from infectious agents | Microbial and viral safety are requested and investigated at different stages of culture | No risk of cross-contamination and variable culture behaviour |
| Raw materials:<br>-Criteria for selection<br>- Feeder layer<br>-Experimental process intermediate | Quality and safety assured by:<br>- Use of compendial materials<br>- Selection of each material according to a specific lot<br>- Maintenance of high clonogenicity, with big colonies of cultured airway epithelial cells<br>- GMP-compliant master cell bank fully tested for viral and other infectious agents<br>- Validated irreversible lethal irradiation protocol<br>- Controls on viability, colony forming efficiency, tissue-specific makers, doubling time, number of cell doublings | Decrease of variability, enhanced safety, and optimization of time for future trial approvals<br>- Minimization of potential contamination of infectious agents<br>- Maximization of TE product performance<br>- Reported absence of 3T3-J2 cells in patients transplanted with other epithelia.<br>- Absence of microchimerism<br>- Reproducibility, minimization of experimental variability, consistency of results, comparability, safety |
| Shelf-life | Stability studies with multiple batches for each reagent/procedure | Optimization of use, optimization of cultures, decreased variability |
| Analysis on cultured human airway epithelial cells | - Long-term replicative senescence, absence of immortalization<br>- Presence of stem cells and proliferation<br>- Presence of all lineages of differentiation<br>- Correct architecture of the tissue<br>- Maintenance of tissue functions | Absence of tumorigenic potential and immortalization events |
| Stem cell content | Measurable and controlled over time | Controlled risk for immortalization, control of long-term tissue regeneration and self-renewal of the tissue |
| Testing | Test procedures validated using the specific cell type | Robustness of procedures, interoperator and intraoperator reproducibility |
| Regulatory oversight | Culture system for epithelia extensively reviewed and approved by the European Medicines Agency for safety of other human epithelia | Multiple experts involved in the evaluations, possibility for quick translation |

Abbreviations: GMP, Good Manufacturing Practice; TE, Tissue Engineering

In terms of infectious hazards, cell culture for advanced therapies needs to avoid cross-contamination by adventitious microbiological agents not covered by traditional donor screening. Numerous viral, bacterial and other pathogenic organisms can grow *in vitro* and were analysed in the culture environment. Given that airway epithelial cells had a high bioburden upon arrival, treatment with amphotericin, gentamycin, penicillin and streptomycin and numerous washing steps were carried out immediately after biopsy collection and before cell extraction.

In specific cases, such as transmissible spongiform encephalopathies ( TSEs), validated tests are not available, although some are emerging that look promising [48].

Risk materials were selected from the European Directorate for the Quality of Medicines & HealthCare (EDQM)-certified materials sourced from TSE-safe geographical areas. Cross-contamination and raw material potency were controlled by using compendial materials, selecting each material according to a specific batch, and testing its ability to maintain high clonogenicity with large colonies of cultured airway epithelial cells. Fundamental biological attributes of cell lines are essential to know; therefore, the cell line used as feeder layer was derived from a GMP-compliant two-tiered master and working cell bank, fully tested for viral and other infectious agents, and validated for irreversible lethal irradiation. All data for expansion, cryopreservation, differentiation, growth beyond passage and the ability to support epithelial regeneration were produced under GMP designed to ensure consistency and high quality in manufacturing [49–51].

Experimental process intermediates and shelf-life were standardized by controls for viability, colony forming efficiency, number of aborted colonies, tissue-specific markers, doubling time, number of cell doublings and by stability studies on multiple batches (see also the following paragraphs and Methods).

The robustness of the expansion process was ensured by measuring long-term replicative senescence, absence of immortalization, presence of stem cells and proliferation, differentiation lineages, correct tissue architecture, maintenance of tissue functions, and a controlled number of stem cells via functional tests. Test procedures were validated using the specific cell type, and regulatory compliance included instrumentation, facilities and records keeping from the initial filling to the final thawing of each vial of cells and production of the airway tissue, with considerable attention paid to training staff to perform each activity reliably.

### Determination of the clonogenic and proliferative potential of human airway primary cultures

Airway epithelial cells were isolated from biopsies of approximately 1, 8 cm^2^ obtained from donors with no history of cancer or systemic disorders.

To assess the clonogenic capacity of different biopsies under the selected culture conditions, a defined number of cells from each sample was plated on lethally irradiated 3T3-J2 cells from the working cell bank prepared according to European Medicinal Agency guidelines [52] and stained 12 days later with rhodamine B. The colony-forming efficiency (CFE) assay of AT (58.87 ± 14%) and AB (66.23 ± 24%) primary cultures revealed an increased number of clonogenic cells compared with those from the biopsy (13.37 ± 11% in AT and 10.87% ± 5% in AB) (**Figure 1A**; **Figure 1B**), in line with previous observations [53]. This increase suggests a selection of proliferating epithelial cells over other nonepithelial cell types (**Figure S1A, B**), supporting the expansion of basal epithelial cells with high proliferative potential in the first passage of culture. In fact, the terminally differentiated cells isolated from the biopsy appear to undergo replicative senescence. The long-term proliferative capacity of cells isolated from different areas of the airways (AT and AB) was subsequently assessed via serial cultivation until replicative senescence. As shown in **Figure 1C**, all mass cultures from AT (n=3, 27-44 passages) and AB (n=3, 24-28 passages) revealed extensive proliferative potential, with 216 ± 52 and 161 ± 16 cell divisions and 20.5 ± 0.5 hours and 19 ± 1.4 hours of duplication for AT and AB epithelial cells, respectively, before senescence. No statistically significant differences in cell doubling or doubling times were observed between the two areas ( **Figure 1D**), indicating that clonogenic cells with a high capacity for cell division (typical stem cells) were evenly distributed in both airway districts.

**Figure 1.**
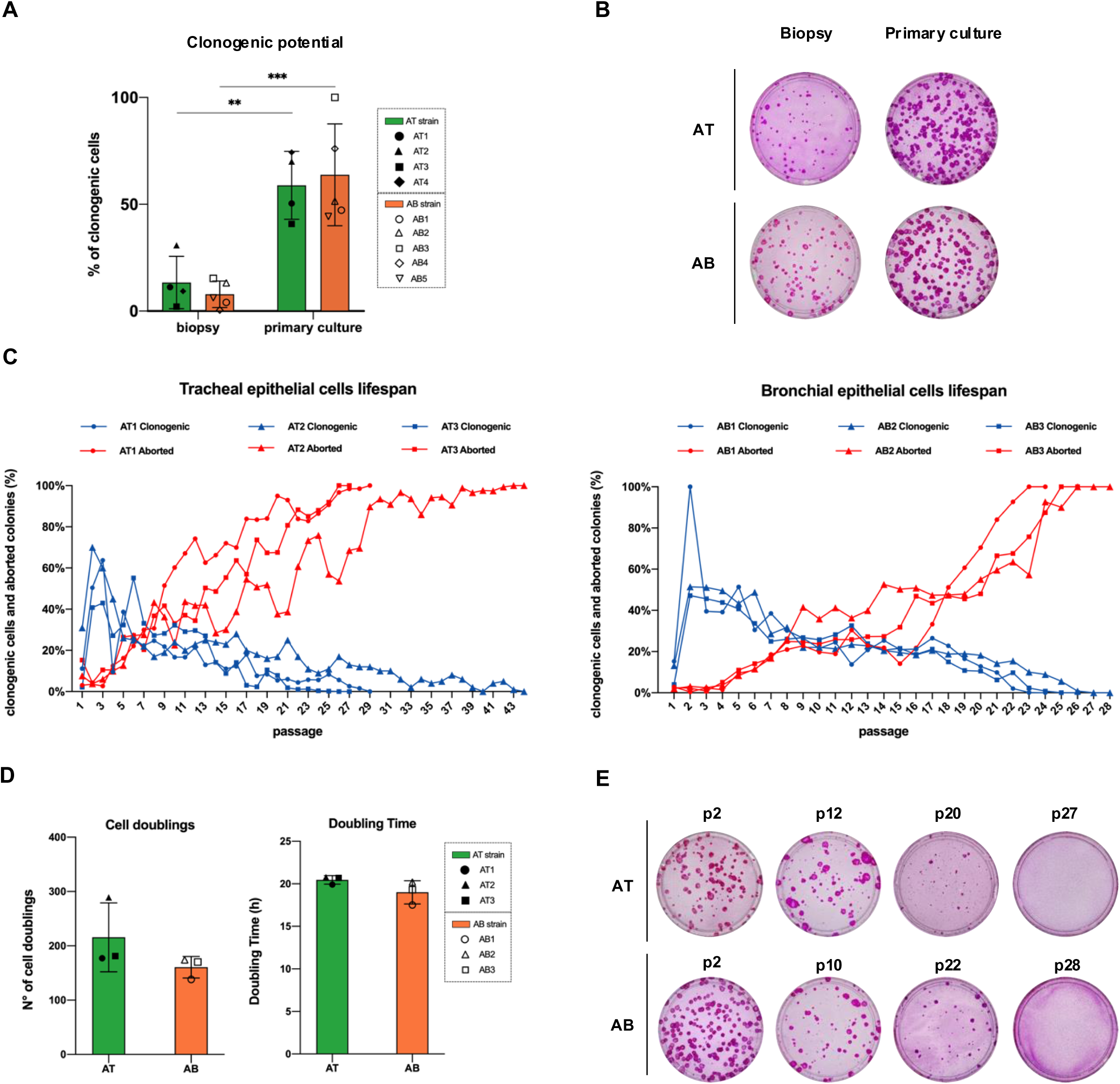
Cell Extraction and Long-term Proliferative Potential of Primary Human Airway Epithelial Cells (A) Comparison of the clonogenicity of cells directly extracted from AT (n = 4) and AB (n = 5) biopsies to that after the first expansion passage. Each independent strain is indicated by a different shape. Unpaired, biparametric, two-tailed t test. (B) Representative images of the CFE assay performed at cell extraction (biopsy, 1000 cells seeded) and after one cell passage (primary culture, 400 cells seeded). (C) Graphs showing quantification of the CFE assay during the AT (left, n = 3) and AB (right, n = 3) lifespans. The blue lines indicate the percentage of clonogenic cells (number of grown colonies/total seeded cell ratio); the red lines indicate the percentage of aborted colonies (aborted colonies/total grown colonies ratio). Independent human strains are indicated by different shapes. (D) Histograms showing the cell doubling (left) and doubling time (right) of AT (n = 3, green bar) and AB (n = 3, orange bar) independent strains, indicated by different shapes. Unpaired, biparametric, two-tailed t test. The values are presented as mean ± SD. ***p < 0.001, **p < 0.01. (E) Representative images of CFE indicator dishes showing the trend of clonogenic and aborted colonies in AT3 and AB2 lifespans. The number of cells seeded in AT3 serial passages was 300 (p2), 500 (p12), 2000 (p20), and 67000 (p27), and the number of cells seeded in AB2 serial passages was 400 (p2), 600 (p10), 700 (p22), and 250000 (p28). p: passages.

No immortalization events were observed, as required in the clinical setting, as shown by p16^ink4A^ (**Figure S1C**). Indeed, after the first passages, a progressive decrease in AT and AB clonogenic cells was accompanied by an increase in terminally differentiated aborted colonies, suggesting physiological progression towards replicative senescence ( **Figure 1C, E**). Differentiated cells were produced throughout the lifespan, excluding basal cell selection, suggesting the occurrence of asymmetric cell divisions.

### Airway cell culture characterization

Epithelia from different parts of the body express cytokeratin (CK) pairs that are unique to each site [54,55]. We therefore analysed the expression of typical epithelial markers during the lifespan of airway cells. The stable expression of CK5 and CK14 within the growing colonies, detected in the native tracheal and bronchial epithelial basal layers (**Figure S2A**), confirms the maintenance of their epithelial basal origin throughout the expansion process ( **Figure 2A; S3A**). In agreement with the *in vivo* results **(Figure S2A)**, the typical marker of pseudostratified epithelia CK18 [56] was expressed in all expanded AT and AB epithelial cells (**Figure S3A**).

**Figure 2.**
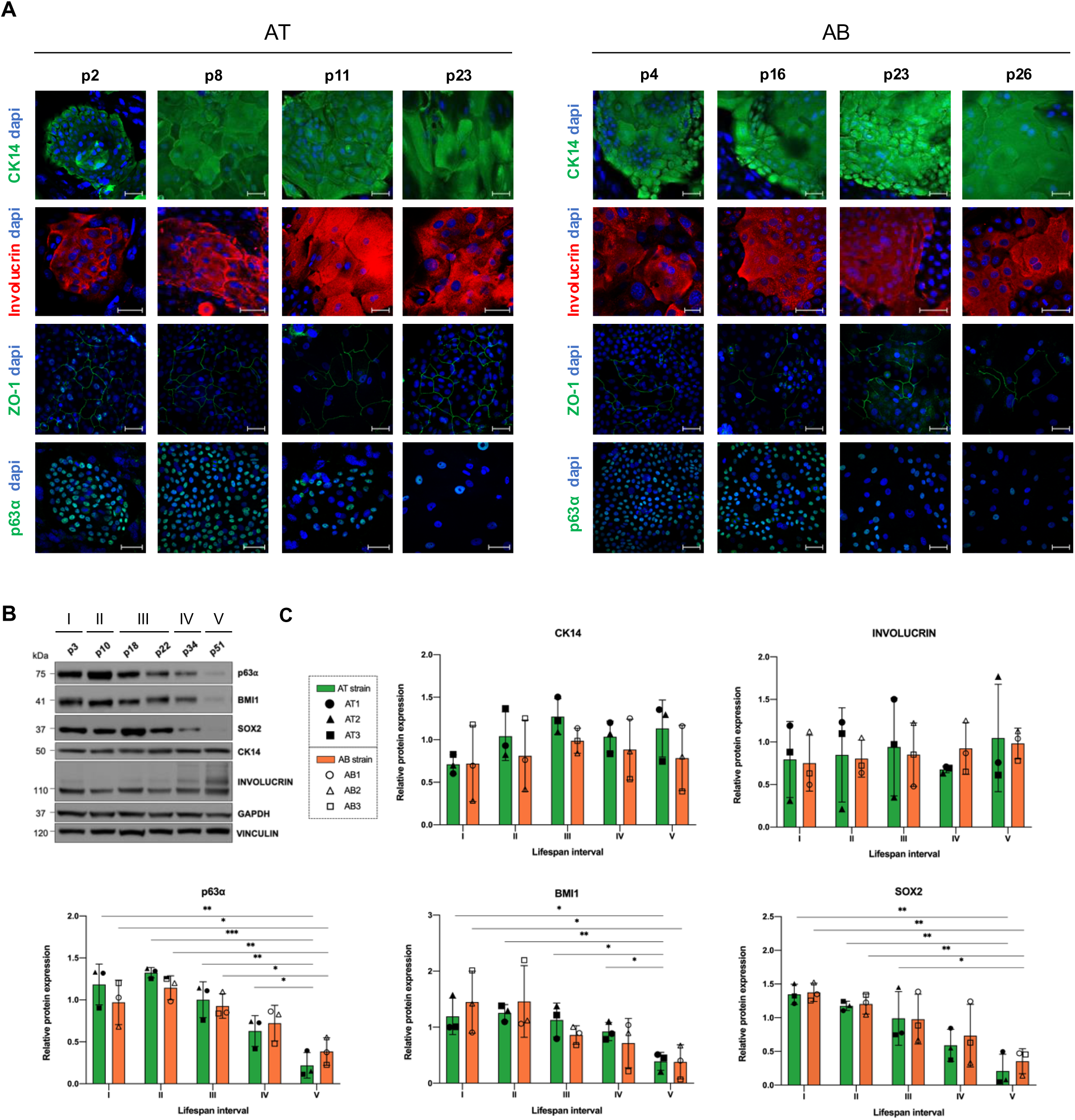
**Characterization of airway epithelial cell cultures** (A) IF staining of AT (left) and AB (right) cultures at four time points during the lifespan of airway epithelial cells (representative images of three AT and three AB primary independent human strains). Scale bar, 50 μm. (B) WB analysis of total cell extracts from six expansion passages (AT2 strain) that represent five consecutive lifespan intervals (I-V), immunostained with the indicated antibodies. The experiment was conducted with n = 3 AT strains (n = 5 technical replicates) and n = 3 AB strains (n = 4 technical replicates). (C) Histograms showing the quantification of the expression levels of CK14, involucrin, p63α, BMI1 (all normalized per GAPDH), and SOX2 (normalized per Vinculin) from the top left to the bottom right. Average and SD of n = 3 AT and n = 3 AB are displayed per time range (see Methods). Independent strains are indicated by different shapes. For multiple pellets analysed within the same interval, the mean expression value was considered. Unpaired, biparametric, two-tailed t test *p < 0.05; ** p < 0.01; and *** p < 0.001.

Detection of the early epithelial differentiation markers involucrin ( **Figure 2A**) and 14-3-3-sigma (or stratifin; **Figure S3A**) confirmed the ability of the cultures to undergo physiological cell differentiation. In particular, involucrin, which was barely detectable in native tissue ( **Figure S2B**), was expressed in most cells of the upper layer of growing colonies ( **Figure 2A**). The expression levels of CK14 and involucrin were quite stable throughout the serial passages of the AT and AB lifespans, with no significant differences detected by WB analysis (**Figure 2B, C**), according to IF analysis.

The cells positively stained for the proliferation marker Ki-67 and the stem/progenitor marker p63α appeared to decrease over serial passages (**Figure 2A, S3A**), consistent with the progressive reduction in clonogenic cells over the lifetime of the culture. These data further confirmed the absence of immortalization events, despite the demonstrated long-term proliferative potential (**Figure 1C**).

In addition, the labelling of the tight junction marker ZO-1 at the cell-cell contact, suggested early regeneration of the epithelial barrier in the colonies and revealed them as well-organized small pieces of tissue (**Figure 2A**).

Notably, a significantly higher expression of three transcription factors (TFs), p63a, BMI1 and SOX2, which are involved in the regulation of stem cell function [57–61], was observed at early AT and AB cell culture time points (time points I and II, enriched in stem and progenitor cells) compared to the end of culture (time point V) (**Figure 2B, C**). The progressive decrease in the expression of these TFs in cells approaching replicative senescence supports the relationship between their high expression and proliferative potential sustained by the stem/progenitor cell population.

### Regenerative and differentiation potential of tracheobronchial epithelial cells

The airway epithelium is populated by several differentiated cells that are essential for maintaining tissue architecture and function. Here, we investigated the production and maintenance of specialized goblet and club cells, which are critical for airway function, throughout the lifespan of airway epithelial cells. Both cell types were detected in all AT and AB cultures, confirming the ability of the culture system to maintain at least two additional tissue-specific lineages (**Figure 3A, B; Figure S3B**). The conditions employed maintained biological variability between independent human strains, as suggested by some fluctuations in the relative number of differentiated cells observed over the life of the cultures (**Figure 3B**). Despite the variability, a specific trend in the development of differentiated cells was observed. Indeed, AB strains presented both club and goblet cells from very early expansion passages, whereas AT strains were able to generate the same cell types later in their lifespan (**Figure 3A, B; Figure S3B**). Intermediate cell types coexpressing typical club and goblet cell markers were also observed in all AT and AB strains (**Figure 3C**). These uteroglobin/MUC5AC- positive hybrid cells may represent a transient state between the two lineages, as previously described [24,62]. Terminal differentiation of airway epithelial cells was then examined in the same culture medium at the air-liquid interface (ALI) to mimic the physiological environment of airway tissue [63–65] (**Figure 3D**). AT and AB cells regenerated a mature, fully differentiated airway epithelia composed of the cell types shown in native tissues (**Figure S2**; **Figure 3E**). Importantly, the presence of properly organized cilia at the apical level demonstrated correct polarization of the regenerated epithelia, whereas secreted mucus supported goblet cell functionality ( **Figure 3E**).

**Figure 3.**
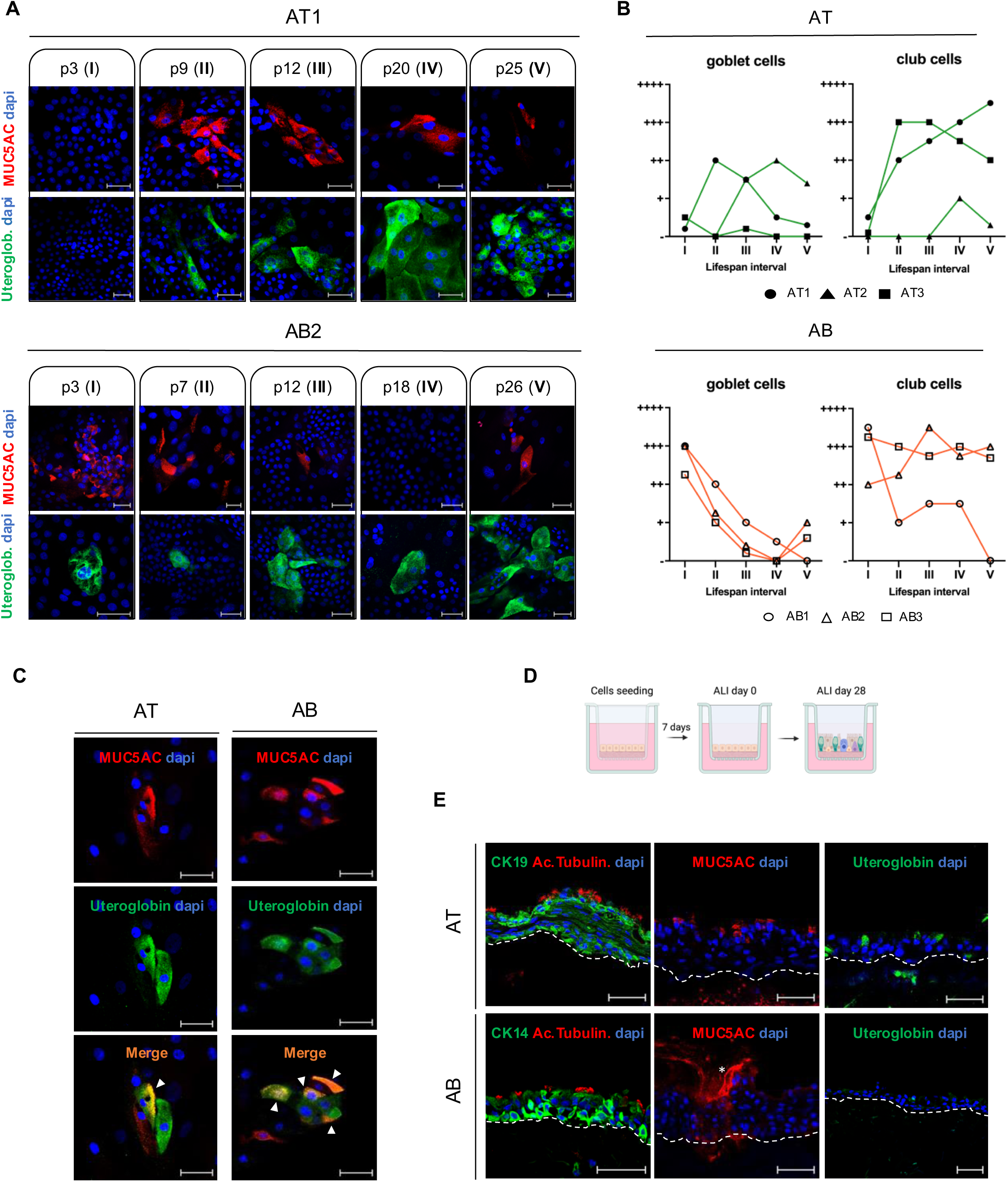
**Regenerative and Differentiation Properties of Human Airway Cultured Cells** (A) IF images showing goblet (MUC5AC-positive) and club (uteroglobin-positive) cells in progressive passages of the AT1 (upper panel) and AB2 (lower panel) lifespans. The same analysis was repeated for the 3 AT and three AB primary strains. Scale bar, 50 μm. (B) Graphical representation of the abundance of goblet and club cells during AT (upper panel) and AB (lower panel) lifespans. Data collected from n = 3 AT and n = 3 AB independent strains are indicated by different shapes. Grading: -, none; +, low; ++, moderate; +++, high; and ++++, very high (see Methods). (C) IF images of MUC5AC (red) and uteroglobin (green) staining in AT and AB cultures. The merged image highlights the presence of double-positive cells coexpressing MUC5AC and Uteroglobin. The white triangles highlight MUC5AC/Uteroglobin double-positive cells. These hybrid cells were observed in n = 3 AT and n = 3 AB-independent strains. Scale bar, 50 μm. (D) Cartoon showing air-liquid interface (ALI) culture protocol. From left to right: cells are seeded onto a de- epithelialized human matrix, cultured for 7 days under submerged conditions, and exposed to ALI for 28 days. (E) Representative images of the epithelium regenerated by AT (n = 3) and AB (n = 3) independent human strains (n = 9 technical replicates). The dotted line indicates the epithelial basal layer; the asterisk indicates the mucous released by goblet cells. Scale bar, 50 μm.

### Basal cell origin of airway culture: single-cell clonal analysis

As shown in **Figures 3A** and **3B**, a variable number of club and goblet cells were progressively observed in AT and AB mass cultures. These data could be explained either by the proliferation of differentiated cells in culture due to the existence of a precursor capable of proliferation w ith restricted differentiation *in vitro* or by the differentiation of multipotent stem cells capable of giving rise to different cell types [45,62,66–68].

We therefore investigated the presence of goblet and club cells in 53 AT and 77 AB clones derived from airway cells. AT and AB individual cells were cultured, and the resulting clones were analysed to determine the generation of the different cell types from each single cell, based on the expression of specific cell markers. All the clones were positive for CK14, confirming their basal cell origin (**Figure 4A, B**). However, club cells were detected in 33% of AT and 72% of AB clones ( **Figure 4A, B**), and goblet cells were detected in 10% of bronchial clones ( **Figure 4A, B**). The greater abundance of the club cell lineage in AB clones and the absence of goblet cells in AT clones confirmed previous findings (**Figure 3B**), suggesting a rapid commitment of bronchial tissue to the two differentiation lineages, whereas tracheal tissue activate the differentiation process later during the regeneration process. Overall, these results suggest that, albeit to different extents and timings, AT and AB clonogenic basal cells are multipotent, as they can give rise to at least epithelial, club and goblet cells (**Figure 3B, 4B**).

**Figure 4.**
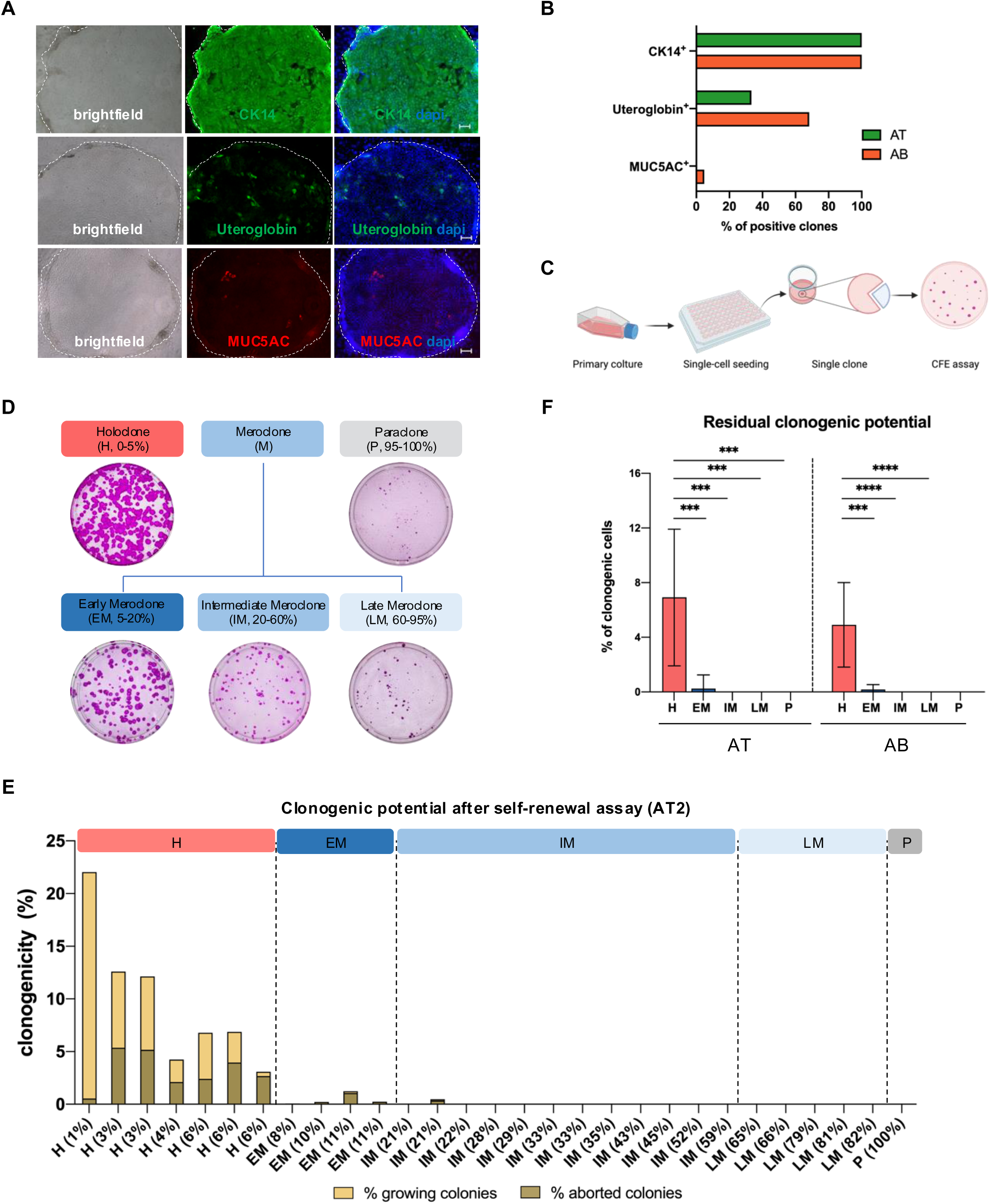
**Basal Cell Heterogeneity and Hierarchy** (A) Representative images of airway clones positive for CK14, Uteroglobin, and MUC5AC. The brightfield image (left) shows the clone morphology. Clones are indicated by white dotted lines. Scale bar, 50 μm. (B) Graph showing the percentage of clones positive for CK14, Uteroglobin, and MUC5AC. CK14 staining was quantified in n = 63 AT and n = 91 AB clones; Uteroglobin staining was quantified in n = 24 AT and n = 29 AB clones; and MUC5AC staining was quantified in n = 48 AT and n = 29 AB clones. (C) Scheme of the clonal analysis procedure. From left to right: subconfluent primary cultures were trypsinized, serially diluted, and inoculated (1 cell per well) into 96-well plates coated with irradiated 3T3-J2 cells. After 7 days of cultivation, part of each clone (one quarter) was plated into an indicator dish to classify the clonal type (see Methods). (D) Representative images of the CFE indicator dishes used to classify the different types of clones based on the percentage of aborted colonies from the total number of grown colonies. Analysis was conducted in n = 3 independent AT strains (384 total AT clones analysed via n = 9 clonal analyses) and in n = 2 AB independent strains (168 total AB clones analysed via n = 4 clonal analyses). (E) Quantification of clonogenicity after the self-renewal assay with AT2 clones. For each clone, the total percentage of grown colonies after 28-30 days of stratification is presented as the sum of the percentage of terminally differentiated colonies (aborted colonies, light brown) and the percentage of colonies that could regenerate the tissue (growing colonies, light yellow). AT2 analysed clones: H, n = 7; EM, n = 4; IM, n = 12; LM, n = 5; P, n = 1. (F) Graphs showing average AT and AB residual clonogenic potential values after stratification of H, EM, IM, LM, and P progeny. The high standard deviation reflects the heterogeneity of H (0-6% of aborted colonies). Analysis was conducted using n = 3 AT strains (H, n = 14; EM, n = 14; IM, n = 29; LM, n = 13 and P, n = 20) and n = 1 AB strain (H, n = 15; EM, n = 4; IM, n = 7; and LM, n = 2). Unpaired, biparametric, two-tailed Welch’s t test was used. The values indicate the means ± SD. ***p < 0.001. ****p < 0.0001.

### Different states of basal cells

As depicted in **Figure 4C**, AT and AB clones (n=384 and n=168, respectively) isolated from AT (n=3) and AB (n=2) epithelial cultures were further characterized to determine their long-term regenerative potential, according to a previously defined classification for the epidermis [44] and other epithelial tissues [45,47,69]. The three main clonal types were holoclones (H, classified by 0- 5% of terminally aborted colonies), meroclones (M, with 5-95% of aborted colonies), and paraclones (P, leading to 95-100% of aborted colonies) with a progressively lower regenerative potential (**Figure 4D**). Our analyses demonstrated the presence of the three clonal types within two airway epithelial tissues and allowed for the first time the isolation of holoclone-forming stem cells from human tracheal and bronchial epithelial cultures.

H accounted for 8.7 ± 2.3% of AT and 23.7 ± 14.4% of AB analysed clones, with inherent variability observed among different strains (**Figure S4A, B**). All M accounted for 77.1± 3.7% of AT and 71.7 ±9.7% of AB clones, and P represented 14.2 ±5.1% of AT and 4.6 ± 4.6% of AB clones ( **Figure S4A**). To better stratify the status of the clonogenic basal cells, the meroclones were further divided into three subgroups: early meroclones (EM, 5-20% of aborted colonies), intermediate meroclones (IM, 20-60% of aborted colonies), and late meroclones (LM, 60-95% of aborted colonies) (**Figure 4D**).

### Holoclone-forming cells maintain self-renewal in the tracheobronchial epithelium

The progeny of each clone was exposed to strong differentiation stimuli (30 days of stratification) to assess the self-renewal ability of the founding cell, maintaining relevant clonogenic and proliferative capacity *in vitro*.

The clones with up to 6% aborted colonies in the CFE were found to retain clonogenic and proliferative potentials after long-term differentiation (**Figure 4E; Figure S4C, D**). These founder clones maintain an original pool of stem cells that can resist continuous differentiation stimuli by self-renewing and dividing into still clonogenic daughter cells, which can regenerate tissue. In addition, all the clones that generated more than 6% of aborted colonies (EM, IM, LM and P) exclusively consisted of TA cells that rapidly underwent replicative senescence. Accordingly, we considered a new threshold (6% of aborted colonies) for defining H in the human tracheobronchial epithelium (**Figure S4C, D**). The difference between the remaining clonogenic ability of H *vs* all other clonal types was statistically significant (**Figure 4F**). Indeed, only a few EM displayed residual growth potential, with a negligible number of colonies, which were almost entirely aborted and could not proliferate further (**Figure 4E, F; Figure S4C, D)**.

### Stem cell marker expression in different tracheobronchial clones

Despite the demonstration that clonal analysis can identify stem cells in tracheobronchial cultures, this assay remains time-consuming and extremely cumbersome [37,70]. Therefore, we evaluated whether airway H could be uniquely identified on the basis of active proliferation, i.e., colony size. No direct correlation was found between AT and AB clone growth (size at 7 days) or clonal category (**Figure 5A**). The same concept applies to doubling time (**Figure S5A, B**), highlighting the inadequacy of these measures to identify stem cell-derived holoclones in airway cultures. Since the expression of p63α, BMI1, and SOX2 was decreased during proliferative senescence of AT and AB cultures (**Figure 2B, C**), we investigated the expression of these TFs within airway clones, as each of these markers is thought to be associated with stem cells in either the airway [58,60,71] or other tissues [57,59].

**Figure 5.**
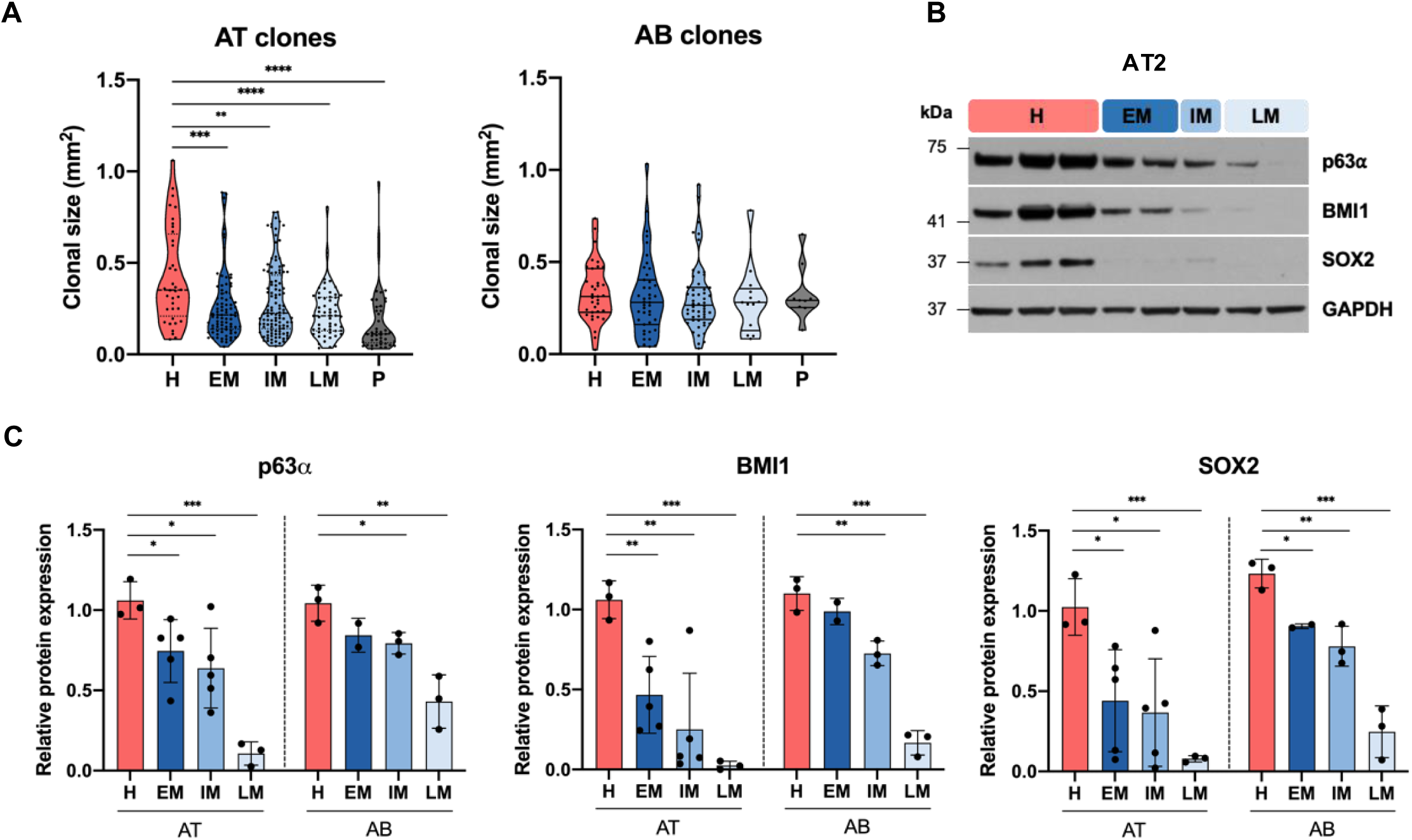
**Morphological and Molecular Characterization of Airway Clones** (A) Violin plot showing the size measured in mm^2^ of AT (left) and AB (right) clones. AT clones: H, n = 36; EM, n = 86; IM, n = 112; LM, n = 56; P, n = 45 belonging to n = 3 independent AT strains; AB clones: H, n = 35; EM, n = 44; IM, n = 52; LM, n = 13; P, n = 9 belonging to n = 2 independent AB strains. Dots represent single clones. Median, first and third quartiles are displayed. (B) WB analysis of total cell extracts from the progeny of AT2 clones (H, n = 3; EM, n = 2; IM, n = 1; LM, n = 2) immunostained with the indicated antibodies (images representative of n = 2 analyses conducted with independent clones). (C) Bar graphs showing the quantification of the expression levels of p63α, BMI1 (normalized to GAPDH), and SOX2 (normalized to Vinculin in AT and to Actin in AB) from left to right. Averages and SD are displayed per clonal category (see Methods). AT2 clones: H, n = 3; EM, n = 5; IM, n = 5; LM, n = 3. AB1 clones: H, n = 3; EM, n = 2; IM, n = 3; LM, n = 3. Due to their very limited residual proliferative potential, the amount of material collected from the progenies of the AT2 and AB1 paraclones was insufficient to carry out a reliable WB analysis. Unpaired, biparametric, two-tailed t test *p < 0.05; ** p < 0.01; *** p < 0.001; and **** p < 0.0001.

Western blot analysis of AT (n=33) and AB (n=11) clones revealed a significant progressive decrease in the expression of these markers from H to LM/P ( **Figure 5B, C; Figure S5C, D**), confirming that p63 a, BMI1 and SOX2 are more abundant in AT and AB epithelial stem cells. Conversely, clones near replicative senescence, although still retaining some proliferative capacity, express extremely low levels of these TFs. These data are consistent with cell behavior during serial culture, where the composition of the mass culture decreases the amount of stem and early progenitor cells (H and EM) to a population composed predominantly of old transiently amplifying and terminally differentiated cells (LM and P) in the latest stage; this phenomenon can be identified as clonal conversion [72–74].

### Differentiation potential of human airway epithelial clones

AT and AB epithelial basal cells showed heterogeneous regenerative capacities. Therefore, to investigate the differentiation potential of stem and transient amplifying cells, we selected 161 airway clones with different proliferative potentials to determine the generation of club, goblet and ciliated cells in the specific state of basal cells ( **Figure 6A, 6C**).

**Figure 6.**
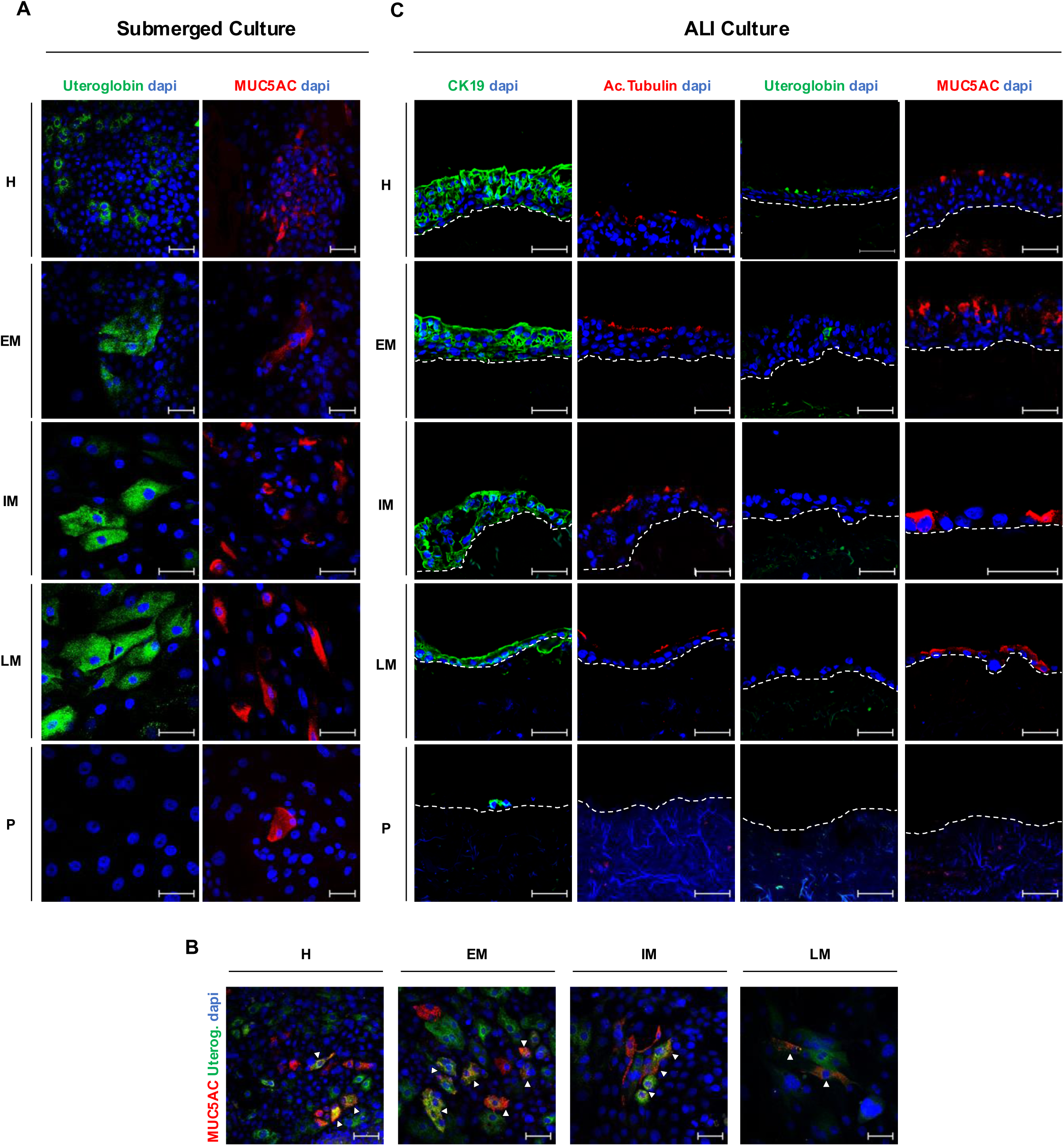
**Differentiation potential of the clones** (A) Representative IF images of the different types of progenies of AT clones cultured under standard submerged conditions. Analysis was conducted using n = 3 AT (H, n = 11; EM, n =10; IM, n = 25; LM, n = 13; P, n = 3) and n = 1 AB strains (H, n = 7; EM, n =6; IM, n =4; LM, n = 4; P, n = 1). The dotted line indicates the epithelial basal layer. (B) Double IF staining of AT and AB epithelial cells revealing coexpression of MUC5AC (red) and Uteroglobin (green) in the progeny of airway H, EM, IM, and LM. White triangles highlight MUC5AC/Uteroglobin-positive cells. (C) Representative IF images of the different types of progenies of AT clones cultured under ALI conditions. Analysis was conducted on n = 3 AT (H, n = 11; EM, n = 19; IM, n = 17; LM, n = 14; P, n = 13) and n = 1 AB strains (H, n = 2; EM, n = 1). Dotted line indicates the epithelial basal layer.

Each clone was cultured under standard and ALI conditions. Standard cultures confirmed the presence of club and goblet cells in the progeny of AT and AB clones, including uteroglobin/MUC5AC-intermediate double-positive cells found in H and all M progeny (**Figure 6A, B**). P-forming cells, the final state of proliferating basal cells, showed limited differentiation potential, as highlighted by the presence of goblet but not club cells in their progeny ( **Figure 6A**). In parallel, the epithelia regenerated from H, EM, IM, and LM ALI cultures were lined on the apical side by ciliated cells (**Figure 6C**). The strong differentiation stimuli imposed by 30 days of ALI culture revealed that H can maintain both regenerative and differentiation potential, confirming its stem cell properties. Conversely, M progeny in ALI conditions were characterized by the absence of club cells within most of the regenerated epithelia (**Figure 6C**). The presence of these cell types in the M cultures exposed to lower differentiation stimuli ( **Figure 6A**) suggests an early exhaustion of this lineage in fully differentiated organotypic cultures. Finally, P was unable to regenerate the epithelium under ALI conditions (**Figure 6C**). Taken together, these results confirm that some, but not all, airway epithelial basal cells can regenerate a fully differentiated tracheobronchial epithelium. Multipotency is not restricted to H stem cells but is maintained in the IM and LM, in contrast to self-renewal. Collectively, these results indicate that H-forming cells are the most powerful and resilient cells, as they can withstand ALI terminal differentiation stimuli and guarantee the maintenance of complex tracheobronchial cellular heterogeneity.

### Functional assays

The efficiency of the culture system in maintaining cells, tissue regeneration, differentiation and functions such as wound healing and the ability to form an effective barrier may not be consistent. Here, the wound healing capacity of airway epithelial cells was tested *in vitro* via a scratch assay. Two culture conditions were compared to test for significant differences in wound healing: the clinical grade medium under investigation (CG) and a standard commercial bronchial epithelial cell growth medium (BEGM). The test involved scratching the tracheal and bronchial epithelia grown under the two culture conditions with a well-defined lesion size and measuring the efficiency and time required for re-epithelialization. Surprisingly, AT and AB epithelia cultured in the CG closed the wound area within 14 hours, whereas their counterparts did not complete the wound healing process after 40 hours (**Figure 7A, B; Supplementary Video 1**). The integrity of the cultured epithelial tissue was tested by analysing the efficiency of its barrier function. This competence was assessed by measuring the transepithelial electrical resistance (TEER) of AT and AB epithelia at different time points in the CG medium under ALI culture environment. TEER measurements every 4-6 days revealed a constant increase in TEER values throughout the culture to a maximum of 421 ± 97 Ω for AT and 361 ± 54 Ω for AB on day 29 of maturation ( **Figure 7C**). These results were consistent with the formation of a dense network of tight junctions observed during the exponential growth phase (**Figure 2A**), confirming the ability of the regenerated epithelium to produce an efficient barrier. Scanning electron microscopy (SEM) highlighted the compactness and polarization of the AT and AB regenerated epithelia, showing large areas densely covered with cilia and shorter microvilli protruding apically from the airway columnar cells ( **Figure 7D**).

**Figure 7.**
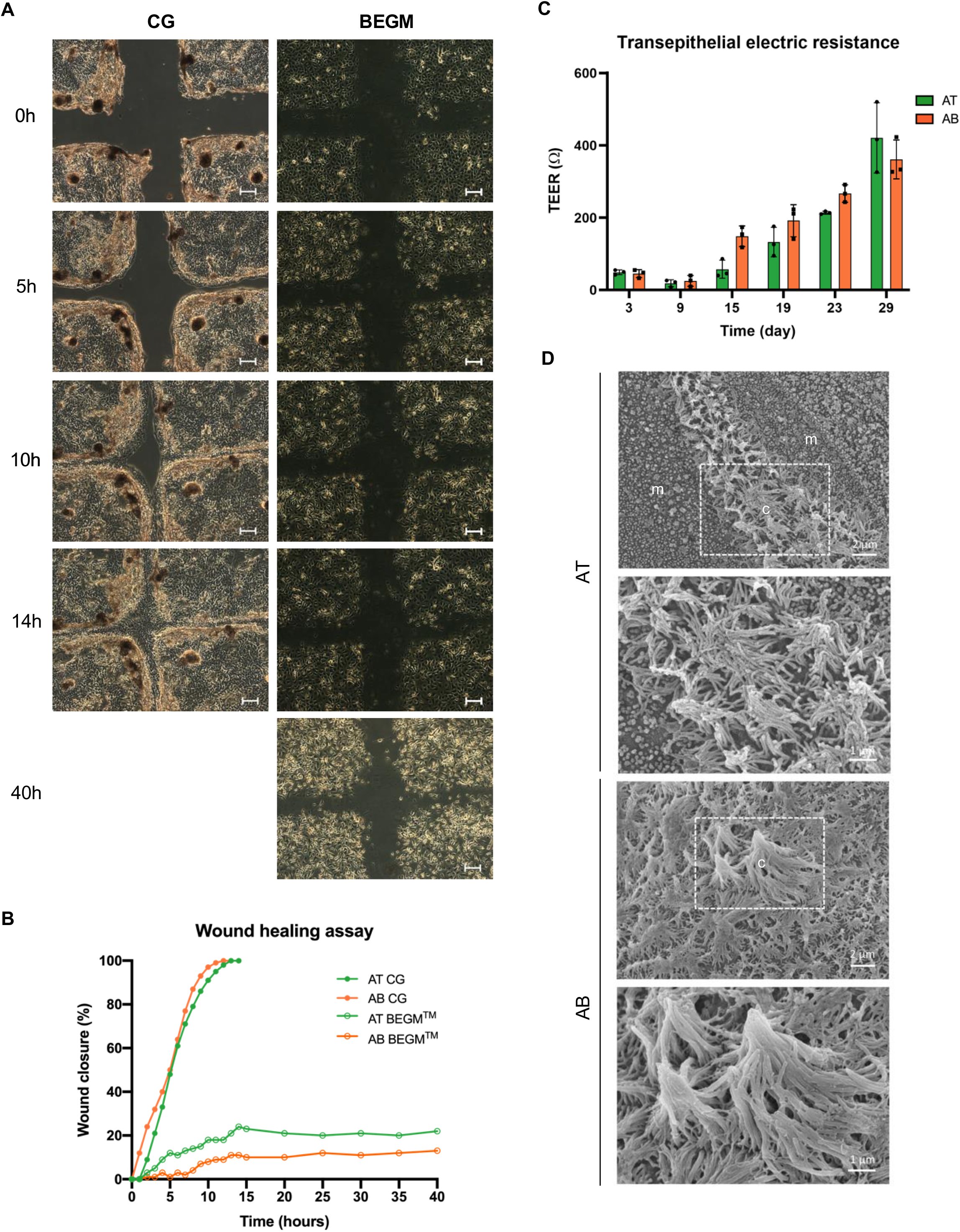
**Assays to Evaluate the Functionality of the Regenerated Airway Epithelium** (A) Representative images of airway epithelia grown in CG (left) or BEGM (right) culture systems at five consecutive time points after the scratch (0 h); images were acquired via live imaging (Cell Observer^®^, Zeiss). The assay was conducted with n = 1 AT and n = 1 AB primary cultures from independent donors. Scale bar, 50 μm. (B) Quantification of the percentage of *in vitro* wound closure in AT and AB epithelia under CG or BEGM culture conditions (see Methods). The wounded area was quantified via AxioVision version 4.8. (C) Graph showing the TEER measurements obtained at six consecutive time points during AT (green bars) and AB (orange bars) ALI cultures. Average. The average and standard deviation are displayed per time point based on three technical replicates of n = 1 AT and n = 1 AB strain. TEER measurements were conducted with an ERS-2 voltmeter (see Methods). (D) Representative SEM images of AT and AB regenerated epithelia showing apical cilia (c) and microvilli (m). The dotted square highlights the area magnified in the corresponding inferior image.

Interestingly, when the epithelium was detached from the support, the superficial ciliated cells rotated in a specific direction, indicating the synchronization of cilia movement and their ability for mucus clearance (**Supplementary Video 2**).

## Discussion

The proposed culture system has been clinically validated for the regeneration of other human epithelial tissues [34,41–43,47]. In this study, all AT and AB epithelial cell cultures were able to maintain a remarkable proliferative capacity and underwent physiological replicative senescence, with a gradual decline in clonogenicity and p63 expression. The absence of immortalization events provided reassurance about the safety of this expansion method, making it less potentially dangerous and more suitable for clinical applications than other approaches. In addition to immortalization procedures [25–27,75,76] that cannot be used in a clinical setting [77,78], techniques based on the inhibition of Rho kinase (ROCK) [32,78–81] or SMAD signalling [82,83] are associated with safety concerns. Although these approaches may prove useful in modelling the airway epithelium *in vitro* [82], inhibition of these pathways can alter epithelial physiology, unbalancing stem cell proliferation, differentiation and cell cycle regulation [84–87]. These changes impair terminal differentiation of airway cells and lead to transient immortalization of the culture, thereby inducing a predominantly stem cell-like phenotype [31].

Here, human airway epithelial cells were found to undergo extensive serial passaging and cell duplication under validated culture conditions that meet the requirements for TE transplantation. The retention of proliferative potential and differentiation capacity has not been thoroughly reported in most unsuccessful and controversial clinical TE attempts [14,88]. We have demonstrated the suitability of clinical-grade expansion conditions to preserve cell heterogeneity and regenerate a fully differentiated airway epithelium capable of fulfilling its functions and acting as an efficient barrier against environmental insults and infections. In particular, the ability to rapidly close a wound, a key feature of TE approaches, was confirmed using this system, but not using a standard available cell culture medium.

Epithelial self-renewal is strictly dependent on functional tissue-specific stem cells [89–94]. More than three decades of clinical applications of cultured human epithelia have shown that the maintenance of stem cells in culture is the key feature determining treatment success or failure [34,35,37,41,95]. Therefore, airway epithelial stem cells have been identified by functional methods such as clonal analysis and self-renewal and characterized by marker expression. Based on our results, the entire clonogenic compartment is composed of basal cells, reinforcing the notion that human tracheal and bronchial epithelia are of basal origin [96–99]. Further studies are needed to verify whether this cultured epithelium, which is composed of basal, club, goblet and ciliated cells, maintains or gives rise to rare airway cell types.

In the present study, we demonstrated that airway basal cells differ in their regenerative capacity and were classified here as H, EM, IM, LM and P, similar but not identical to what has previously been demonstrated in other human epithelia [44,45,47,69]. This result highlights the heterogeneity of the airway basal compartment and suggests that not all basal cells can equally guarantee the complexity of tissue renewal and long-term maintenance. Most importantly, tracheal and bronchial H-forming cells have specific characteristics of epithelial stem cells, and, similar to observations in other epithelia [35,42], human airway H-forming cells, but not M or P, can self-renew and withstand strong differentiation stimuli. The EM, IM, LM and P, which are transient progenitors, progressively lose their proliferative potential and thus contribute to tissue regeneration for a limited time. In fact, all H showed self-renewal capacity; however, the variable clonogenic potential after stratification contributes to the ongoing debate as to whether all H-forming cells are identical or whether there is a hierarchical relationship between them [35]. H must be present, and an adequate amount is required to achieve successful long-term tissue regeneration and therapies [37,41]. Clonal analysis, which is the gold standard method for identifying stem cell function in many epithelia, revealed the preservation of airway H during expansion and can be applied during the validation of the most critical steps of the TE procedure. Therefore, the quantification of stem cells within an epithelial culture and the maintenance of a sufficient number of stem cells are of paramount importance [39,70] in controlling the manufacturing process, in addition to proliferation and differentiation.

Clonal analysis, which is important in validation procedures, cannot be used for routine in-process control because it is cumbersome and time-consuming; moreover, analyses of clonal size and growth rate do not identify airway stem cells (H), as reported for other human epithelia [42,44,45,70,100]. The combined evaluation of p63α, BMI1 and SOX2 TFs provided more substantial results. Consistent with the airway epithelial lifespan data, these TFs were significantly differentially expressed between the stem cell clones (H) and transient progenitor cells (all M and P), ultimately strengthening their correlation with stem cell properties. This observation seems consistent with data showing the indispensable role of p63 in maintaining the proliferative potential of stem cells in adult stratified epithelia [73,101,102], including the airway epithelium. Accordingly, p63 ablation in mice results in offspring lacking stratified squamous epithelia and a complete absence of basal airway cells [60,101]. Similarly, BMI1 plays a critical role in maintaining the self-renewal of several adult stem cells, such as hematopoietic [103], neural [104] and bronchioalveolar stem cells [58]. Finally, SOX2 is essential not only for the self-renewal and pluripotency of embryonic stem cells [59,61] but also for lung development [71], where it plays a key role in airway epithelial proliferation and lineage specification [105,106]. Notably, SOX2 overexpression in the airway epithelium led to an increase in the number of p63-positive cells, suggesting their synergy in maintaining and upregulating basal stem cells [71]. The described findings of differential p63 and BMI1 expression in airway clones are consistent with previously reported patterns in epidermal, corneal, oral mucosa and urethral clones [39,47,57,73]. The robust association between the number of p63-rich holoclones and clinical success confirmed p63 as a potency marker [37,41] in limbal cultures and led to the approval of the first stem cell-based ATMP by the European Medicines Agency [33].

This differential expression of TFs in airway clones could be used to quantify holoclone-forming stem cells in airway cultures designed for cell therapy and TE approaches. The critical specifications for controlling product performance/potency can advance airway TE toward standardized, safe and effective therapies.

Notably, in this study, we also demonstrated the preservation of airway epithelial heterogeneity. Indeed, both airway stem (H) and progenitor (M) cells can regenerate a fully specialized airway epithelium composed of the primary cell types that characterize the native tissue. These findings complement the observations of Kumar et al. [97] and highlight the potential of individual tracheal epithelial basal cells to differentiate into club cells in addition to ciliated and goblet cells. Furthermore, this multipotency was maintained until late transient amplifying cells (LM), but was absent in P (**Figure 8**), suggesting the loss of multipotency and irreversibility of clonal conversion. Thus, stem and progenitor cells are responsible for replenishing differentiated cell lineages during physiological airway epithelial turnover and wound healing. A similar mechanism has been described in the conjunctival epithelium, where goblet cells are derived from both stem and progenitor cells [45], and in the thymus, where the stem cell refractive index and progenitor stratified clones share a common differentiation potential [69]. Interestingly, airway P-basal cells, which are still capable of limited proliferation, lose some of their multipotency and generate only epithelial and goblet cells, suggesting a point of no return for the generation of specific differentiation lineages during clonal conversion with the need for sufficient proliferative potential to generate club and ciliated cells [107,108]. Club cells appear earlier than goblet cells in basal clones and are absent in M cultures under strong differentiation stimuli such as ALI.

**Figure 8.**
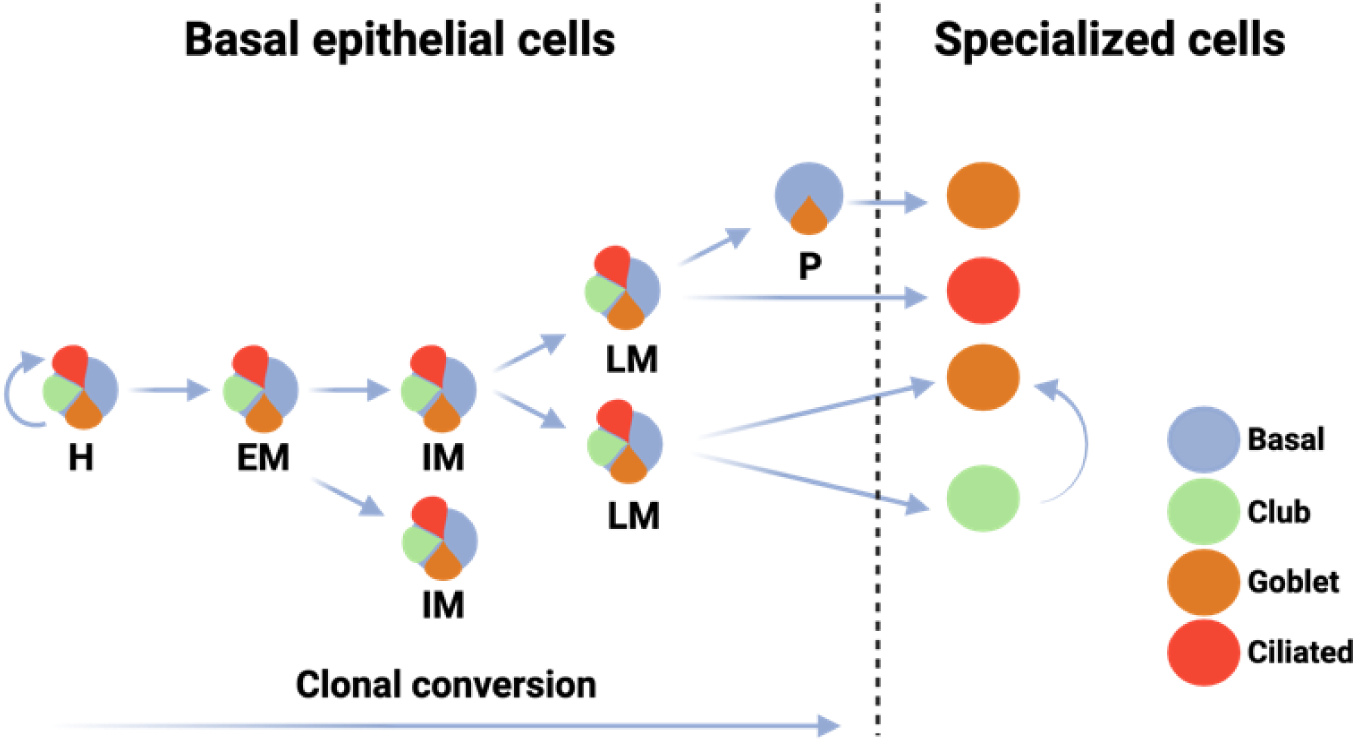
**Airway clone renewal and differentiation potential.** Cartoon showing a model of *in vitro* human airway epithelium renewal and differentiation potential of clones. H, EM, IM, and LM basal clones display multipotency in differentiating into club, goblet, and ciliated cells, whereas paraclones can only differentiate into goblet cells. Among specialized cell types, club cells may differentiate into goblet cells.

The presence of uteroglobin/MUC5AC-positive cells, also reported by other groups [24,82,109,110], suggests a progenitor role for club cells, which can transform into other specialized cell types in the airway epithelium, including goblet cells [109,111,112]. This observation is consistent with findings in animal models generated via injury and lineage tracing experiments [24,107,113] and with hypotheses derived from single-cell RNA sequencing analysis in humans [62]. Further studies will be required to confirm the potential of human club and goblet cells to directly differentiate into ciliated cells, as suggested by some authors [62,68].

## Conclusions

In conclusion, our results suggest that autologous human airway epithelial cells are a suitable cell source for TE approaches under appropriate culture conditions. Inadequate expansion methods may accelerate airway clonal conversion, leading to premature exhaustion of the stem cell population, as demonstrated in epidermal keratinocytes [74] and the loss of some differentiation lineages. In such scenarios, the culture may not result in stable regeneration of the airway epithelium after transplantation, ultimately leading to failure of TE-based treatment with serious consequences for patient health. In-process controls are required throughout the manufacturing process to ensure the quality of the airway epithelium in bioengineered grafts. Here, we propose validations and quality controls, including (i) conditions for airway biopsy collection and cell extraction methods, which have a significant impact on cell culture performance and overall reproducibility of results; (ii) methods for characterizing cultured airway cells to meet safety and identity requirements; (iii) test the maintenance of long-term proliferative potential and generation of all differentiation lineages during the validation of reagents for the TE process, and (iv) assessment of an adequate number of stem cells to ensure long-term graft regeneration. This latter control should be performed via functional assays, such as clonal analysis or other specific assays.

Insights gained into tissue-specific biology and epithelial regeneration under *in vitro* expansion can help overcome critical hurdles in airway TE and promote safe and effective translational research.

## List of abbreviations

TE: Tissue engineering; AT: Airway Tracheal; AB: Airway Bronchial; CG: Clinical-Grade; GMP: Good Manufacturing Practice; FL: Feeder Layer; CFE: Colony-forming efficiency; H: Holoclones; EM: Early Meroclones; IM: Intermediate Meroclones; LM: Late Meroclones; P: Paraclones; SEM: Scanning Electron Microscopy; BEGM: Bronchial Epithelial Growth Medium; TEER: Trans- Epithelial Electrical Resistance; ALI: Air-liquid Interface; SD: Standard Deviation; ATMP: Advanced Therapy Medicinal Product; TSE: transmissible spongiform encephalopathies; EDQM: European Directorate for the Quality of Medicines & HealthCare; CK: Cytokeratin; WB: Western Blot; IF: Immunofluorescence; TFs: Transcription Factors.

## Declarations

### Ethical approval and consent to participate

All specimens were obtained in accordance with the tenets of the Declaration of Helsinki and anonymized. Human tracheal and bronchial samples were collected from n=5 males and n=5 female donors aged between 38 and 76 years. The donors provided informed consent for the use of their biological material for the present study, and ethical committee approval was obtained from all involved centres “Arcispedale S. Maria Nuova di Reggio Emilia”, IRCCS, Reggio Emilia, Italy, (title “Pre-clinical study aimed at characterizing human respiratory epithelial stem cells, their differentiation pathways and potential use as an in vitro model for toxicity studies and tissue engineering”; “Comitato Etico Area Vasta Emilia Nord”, Protocol N. 2019/0014725; date of approval 05/02/2019 ), and the “Fondazione Policlinico Universitario A. Gemelli”, IRCCS, Rome, Italy (title “Pre-clinical study aimed at characterizing human respiratory epithelial stem cells, their differentiation pathways and potential use as an in vitro model for toxicity studies and tissue engineering”; “Comitato Etico Territoriale Lazio Area 3” N. 0008968/21; date of approval 10/03/2021). Human skin samples were obtained from “Azienda Ospedaliero-Universitaria di Modena”, IRCSS, Modena, Italy” from healthy living donors upon informed consent and compliance with Italian regulations (title “Human epithelial stem cells: characterization and development of clinical applications in Regenerative Medicine”; Comitato Etico dell’Area Vasta Emilia Nord, number 178/09; date of approval 29/09/2009).

### Consent for publication

All the authors provided consent for publication.

### Availability of data and materials

Data sharing is not applicable to this article as no data sets were generated or analyzed during the current study.

### Author Contributions

Conceptualization: G.P., D.A., V.G.G.; formal analysis: V.G.G., D.A., GG. ; investigation: D.A., V.G.G., G.G., C.C., A.M., F.L., F.B.; human samples collection: F.L.C., J.E.; writing-original draft preparation: D.A., V.G.G., G.G.; writing-review and editing: G.P., D.A., V.G.G., G.G., C.C., A.M., F.L., F.B., D.Q., J.E., F.L.C.; supervision: G.P.; funding acquisition: G.P. All the authors have read and agreed to the published version of the manuscript.

## Acknowledgements

We would like to thank Anna Ribbene and Sonia Carulli for their help with cell culture. The authors declare that they have not use AI-generated work in this manuscript.

## Supplementary information

Supplementary video 1. *In vitro* wound closure of AT cells

Wound healing assay performed on a confluent monolayer of AT cells grown in CG or BEGM™ culture medium. Images were taken through live-imaging (Cell Observer®, Zeiss) every 10 min for 14 hours and then every 300 min for up to 40 hours.

Supplementary video 2. Ciliated cell functionality

Video showing a cluster of ciliated cells detached from the support after 30 days of ALI culture moving and spinning within the cell culture plate.

**Figure S1. Cell composition of primary cultures and expression of senescence marker during the lifespans**

(A) Representative images of IF analysis conducted on cytocentrifuge samples derived from airway primary cultures stained for epithelial (CK14, red) and feeder layer (mouse-specific Vimentin, green) markers. Scale bar, 50 μm.

(B) Quantification of the cell composition tested in an airway primary culture (AB2). The percentage of CK14+ epithelial cells was calculated as 91,4%, the vimentin-positive 3T3 F/L cells accounted for 6,1% of the total cell composition, while only a few CK14- and vimentin- cells (2,5%) were still present after the first passage of culture.

(C) WB analysis of total cell extracts from serial passages representative of n=3 AT and n=3 AB cultures lifespan immunostained for p16 senescence marker. p16 expression levels were normalized to those of Vinculin.

**Figure S2. Immunofluorescence Analysis of Human Tracheal and Bronchial Samples**

(A-B) Representative images of *in vivo* sections of tracheal (n = 3) and bronchial (n = 3) human specimens immunostained for cytokeratins (CK5, CK14, CK18, CK19), stem-progenitor (p63α) and proliferative (Ki-67) cell markers, airway-specific cell markers (Acetylated Tubulin; MUC5AC, Uteroglobin), and epithelial differentiation (Involucrin) and integrity (ZO-1) markers. Scale bar, 50 μm. AT: Airway Trachea; AB: Airway Bronchus; Ac. Tubulin: Acetylated Tubulin.

**Figure S3. Airway Cell Culture Characterization and Differentiation Potential**

(A) IF staining of AT (left) and AB (right) cultures at four time points during the lifespan of airway epithelial cells. Images represent three AT and three AB primary human-independent strains. Scale bar, 50 μm.

(B) IF images showing goblet (MUC5AC-positive) and club (uteroglobin-positive) cells in the progressive passages of AT2 and AT3 (upper panel), and AB1 and AB3 (lower panel) cultures’ lifespan. Scale bar, 50 μm.

**Figure S4. Self-renewal Assay of the AT and AB Strains**

(A) Table reporting the mean ± SD of the relative abundance of H, M and P in AT (n=3) and AB (n=2) strains.

(B) Distribution of the clonal types within the analysed AT (n=3) and AB (n=2) strains.

(C) Representative images of the CFE indicator dishes used to classify the different types of clones at t0 and evaluate the residual clonogenicity of the progeny subjected to a strong differentiation stimulus (stratification) at 28 days after cultivation (t1). As clones containing up to 6% aborted colonies at t0 retained significant clonogenic potential at t1, they were classified as H.

(D) Quantification of the percentage of colonies grown in the CFE indicator dishes. From top to bottom: AT1, AT3, and AB1 clones at the end of the self-renewal assay (t1). For each clone, the total percentage of grown colonies after stratification is shown as the sum of the percentage of aborted colonies (light brown) and the percentage of colonies that can regenerate the tissue (growing colonies, light yellow). Analysis was conducted using n = 19 AT1 clones (H, n = 1; IM, n = 7; LM, n = 4; P, n = 8); on n =41 AT3 clones (H, n = 6; EM, n = 9; IM, n = 11; LM, n = 4; P, n = 11); and n = 28 AB1 clones (H, n = 15; EM, n = 4; IM, n = 7; LM, n = 2).

**Figure S5. Growth Rate and Molecular Characterization of Airway Clones**

(A) Floating bars showing the doubling time (hours) of AT1 clones (H, n = 8; EM, n = 19; IM, n = 14; LM, n = 7; P, n = 9); AT2 clones (H, n = 8; EM, n = 13; IM, n = 9; LM, n = 4); and AT3 clones (H, n = 7; EM, n = 10; IM, n = 6; LM, n = 1). A progressive increase in doubling time was observed from H to P. Unpaired, biparametric, two-tailed t test *p < 0.05; ** p < 0.01; *** p < 0.001 and **** p < 0.0001.

(B) Floating bars indicating the doubling time (hours) of AB1 clones (H, n = 13; EM, n = 18; IM, n = 7; LM, n = 3) and Normal Human Bronchial Epithelial (NHBE) clones (H, n = 3; EM, n = 14; IM, n = 14; LM, n = 1; P, n = 1). Unpaired, biparametric, two-tailed t test *p < 0.05; ** p < 0.01; *** p < 0.001 and **** p < 0.0001.

(C) WB analysis of total cell extracts from the progeny of AB1 clones (H, n = 3; EM, n = 2; IM, n = 3; LM, n = 3) immunostained with the indicated antibodies.

(D) Images and quantification based on WB analysis of total cell extracts from the progeny of AT1 (top panel: H, n = 2; EM, n = 1; IM, n = 2; LM, n = 1; P, n =2) and AT3 clones (bottom panel: H, n = 1; EM, n = 2; IM, n = 3; LM, n = 1; P, n = 2) immunostained with the indicated antibodies. Graphs show the averages and SD of each clonal category.

**Supplementary Table 1.**
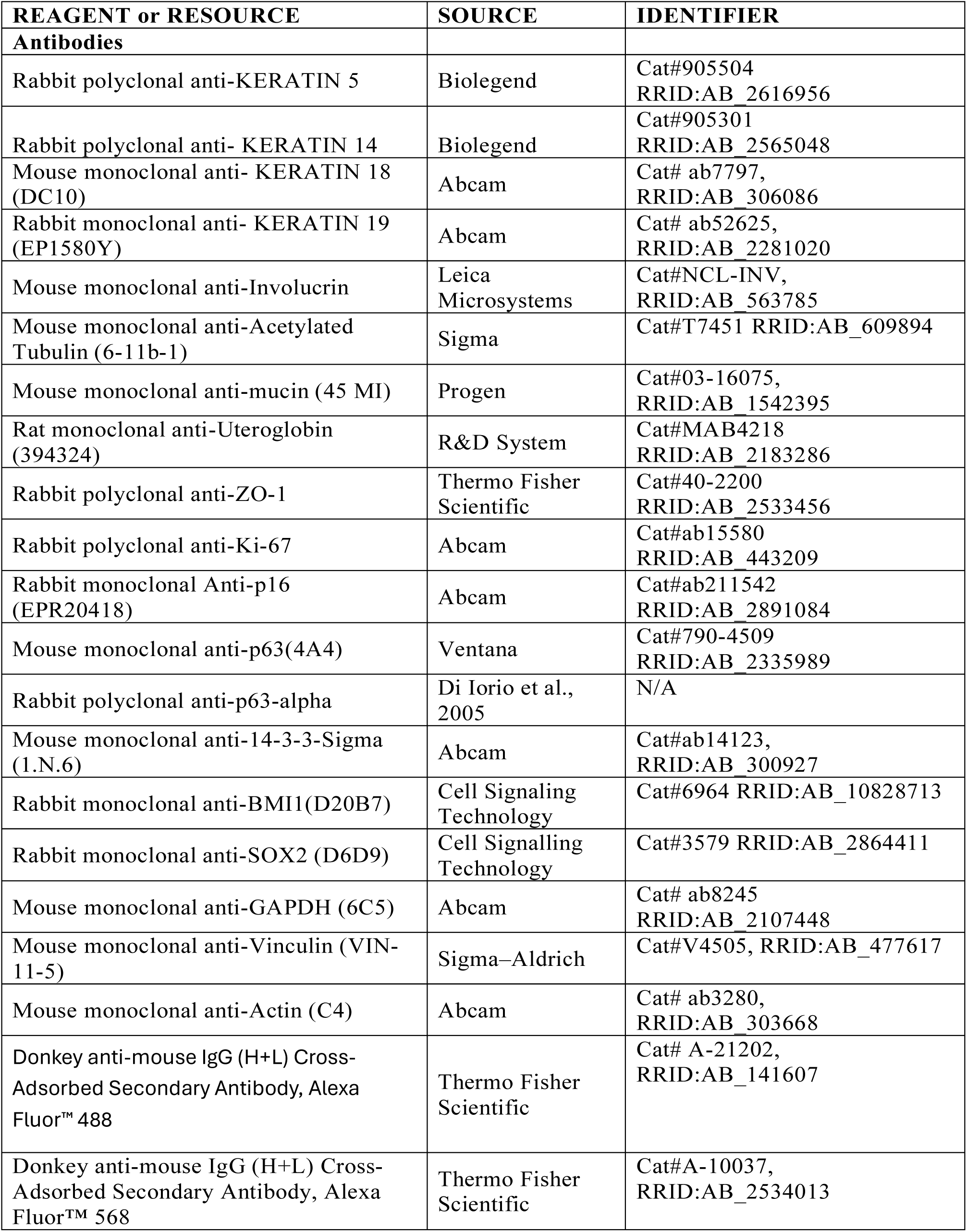
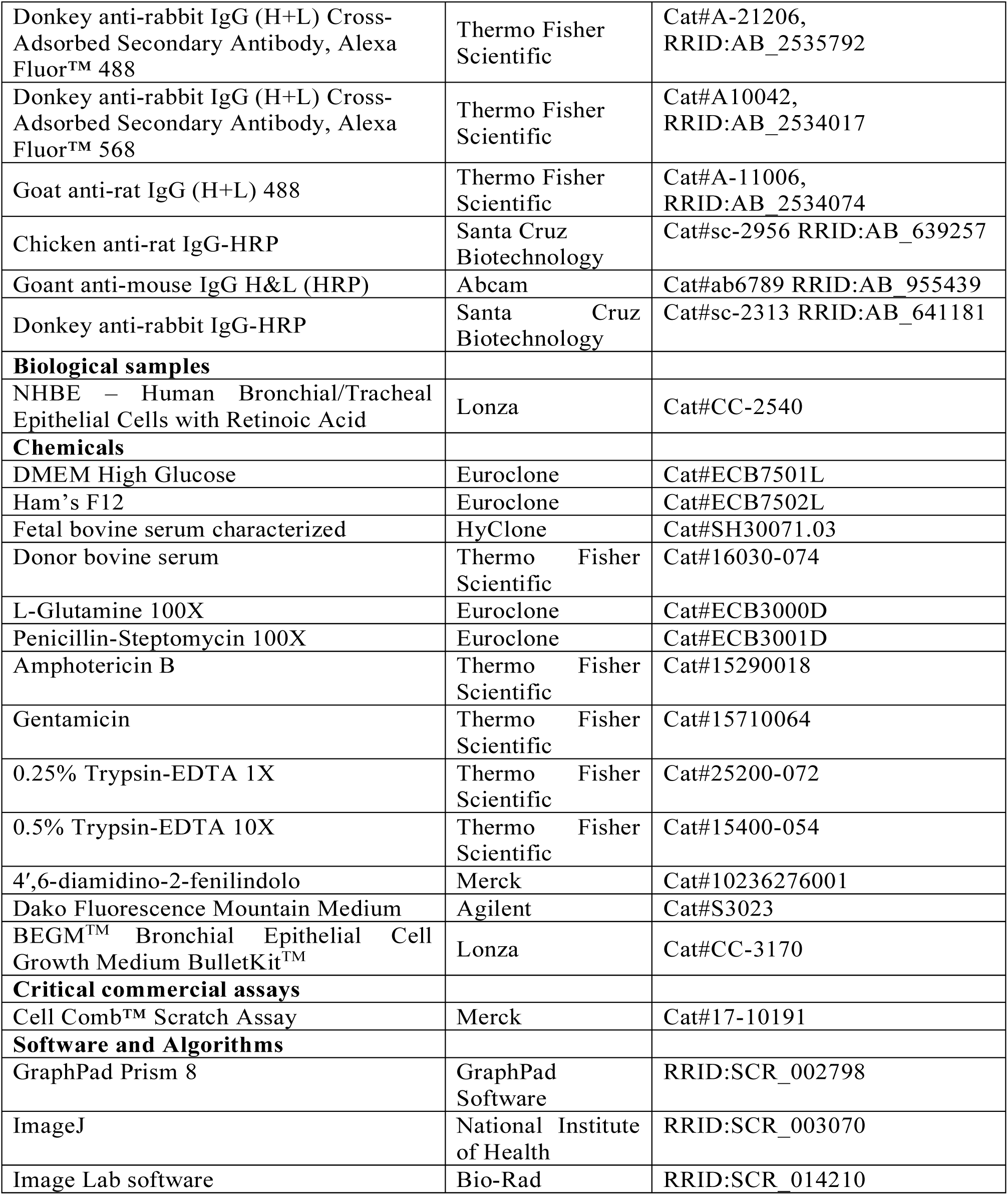
List of reagents and resources employed.

| REAGENT or RESOURCE | SOURCE | IDENTIFIER |
| --- | --- | --- |
| **Antibodies** |  |  |
| Rabbit polyclonal anti-KERATIN 5 | Biolegend | Cat#905504<br>RRID:AB_2616956 |
| Rabbit polyclonal anti- KERATIN 14 | Biolegend | Cat#905301<br>RRID:AB_2565048 |
| Mouse monoclonal anti- KERATIN 18 (DC10) | Abcam | Cat# ab7797,<br>RRID:AB_306086 |
| Rabbit monoclonal anti- KERATIN 19 (EP1580Y) | Abcam | Cat# ab52625,<br>RRID:AB_2281020 |
| Mouse monoclonal anti-Involucrin | Leica Microsystems | Cat#NCL-INV,<br>RRID:AB_563785 |
| Mouse monoclonal anti-Acetylated Tubulin (6-11b-1) | Sigma | Cat#T7451 RRID:AB_609894 |
| Mouse monoclonal anti-mucin (45 MI) | Progen | Cat#03-16075,<br>RRID:AB_1542395 |
| Rat monoclonal anti-Uteroglobulin (394324) | R&D System | Cat#MAB4218<br>RRID:AB_2183286 |
| Rabbit polyclonal anti-ZO-1 | Thermo Fisher Scientific | Cat#40-2200<br>RRID:AB_2533456 |
| Rabbit polyclonal anti-Ki-67 | Abcam | Cat#ab15580<br>RRID:AB_443209 |
| Rabbit monoclonal Anti-p16 (EPR20418) | Abcam | Cat#ab211542<br>RRID:AB_2891084 |
| Mouse monoclonal anti-p63(4A4) | Ventana | Cat#790-4509<br>RRID:AB_2335989 |
| Rabbit polyclonal anti-p63-alpha | Di Iorio et al., 2005 | N/A |
| Mouse monoclonal anti-14-3-3-Sigma (1.N.6) | Abcam | Cat#ab14123,<br>RRID:AB_300927 |
| Rabbit monoclonal anti-BMI1(D20B7) | Cell Signaling Technology | Cat#6964 RRID:AB_10828713 |
| Rabbit monoclonal anti-SOX2 (D6D9) | Cell Signalling Technology | Cat#3579 RRID:AB_2864411 |
| Mouse monoclonal anti-GAPDH (6C5) | Abcam | Cat# ab8245<br>RRID:AB_2107448 |
| Mouse monoclonal anti-Vinculin (VIN-11-5) | Sigma–Aldrich | Cat#V4505, RRID:AB_477617 |
| Mouse monoclonal anti-Actin (C4) | Abcam | Cat# ab3280,<br>RRID:AB_303668 |
| Donkey anti-mouse IgG (H+L) Cross-Adsorbed Secondary Antibody, Alexa Fluor™ 488 | Thermo Fisher Scientific | Cat# A-21202,<br>RRID:AB_141607 |
| Donkey anti-mouse IgG (H+L) Cross-Adsorbed Secondary Antibody, Alexa Fluor™ 568 | Thermo Fisher Scientific | Cat#A-10037,<br>RRID:AB_2534013 |
| Donkey anti-rabbit IgG (H+L) Cross-Adsorbed Secondary Antibody, Alexa Fluor™ 488 | Thermo Fisher Scientific | Cat#A-21206, RRID:AB_2535792 |
| Donkey anti-rabbit IgG (H+L) Cross-Adsorbed Secondary Antibody, Alexa Fluor™ 568 | Thermo Fisher Scientific | Cat#A10042, RRID:AB_2534017 |
| Goat anti-rat IgG (H+L) 488 | Thermo Fisher Scientific | Cat#A-11006, RRID:AB_2534074 |
| Chicken anti-rat IgG-HRP | Santa Cruz Biotechnology | Cat#sc-2956 RRID:AB_639257 |
| Goat anti-mouse IgG H&L (HRP) | Abcam | Cat#ab6789 RRID:AB_955439 |
| Donkey anti-rabbit IgG-HRP | Santa Cruz Biotechnology | Cat#sc-2313 RRID:AB_641181 |
| **Biological samples** |  |  |
| NHBE – Human Bronchial/Tracheal Epithelial Cells with Retinoic Acid | Lonza | Cat#CC-2540 |
| **Chemicals** |  |  |
| DMEM High Glucose | Euroclone | Cat#ECB7501L |
| Ham's F12 | Euroclone | Cat#ECB7502L |
| Fetal bovine serum characterized | HyClone | Cat#SH30071.03 |
| Donor bovine serum | Thermo Fisher Scientific | Cat#16030-074 |
| L-Glutamine 100X | Euroclone | Cat#ECB3000D |
| Penicillin-Streptomycin 100X | Euroclone | Cat#ECB3001D |
| Amphotericin B | Thermo Fisher Scientific | Cat#15290018 |
| Gentamicin | Thermo Fisher Scientific | Cat#15710064 |
| 0.25% Trypsin-EDTA 1X | Thermo Fisher Scientific | Cat#25200-072 |
| 0.5% Trypsin-EDTA 10X | Thermo Fisher Scientific | Cat#15400-054 |
| 4',6-diamidino-2-phenylindole | Merck | Cat#10236276001 |
| Dako Fluorescence Mounting Medium | Agilent | Cat#S3023 |
| BEGM™ Bronchial Epithelial Cell Growth Medium BulletKit™ | Lonza | Cat#CC-3170 |
| **Critical commercial assays** |  |  |
| Cell Comb™ Scratch Assay | Merck | Cat#17-10191 |
| **Software and Algorithms** |  |  |
| GraphPad Prism 8 | GraphPad Software | RRID:SCR_002798 |
| ImageJ | National Institute of Health | RRID:SCR_003070 |
| Image Lab software | Bio-Rad | RRID:SCR_014210 |

