## Supplementary figures and images for "Towards Clinical-Grade Bioengineered Airways: A Study on Stem Cell Renewal and Epithelial Differentiation"

### supplemental Figures

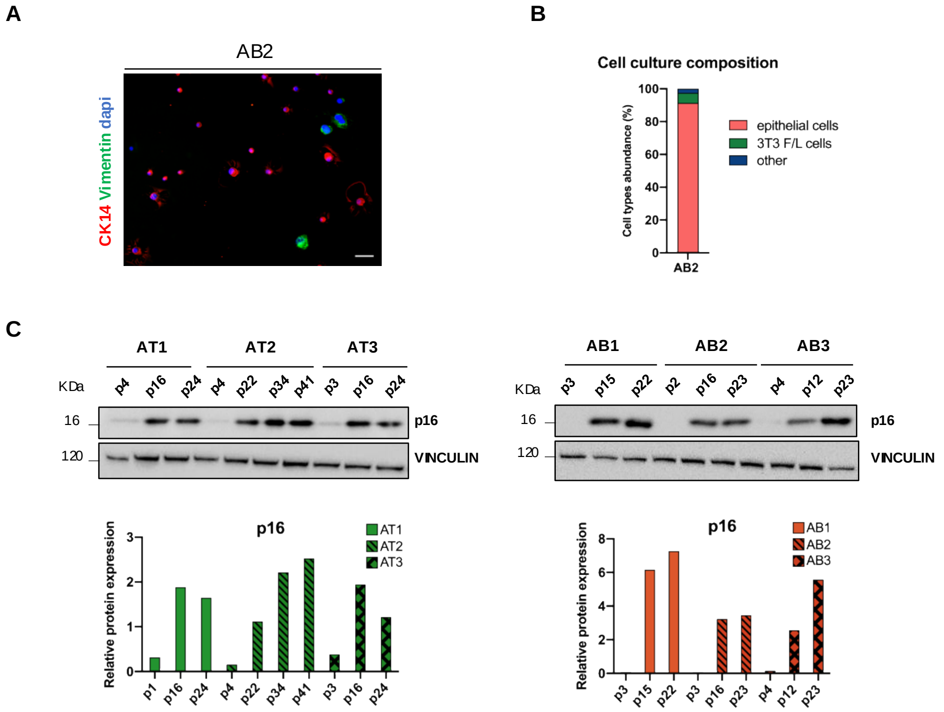

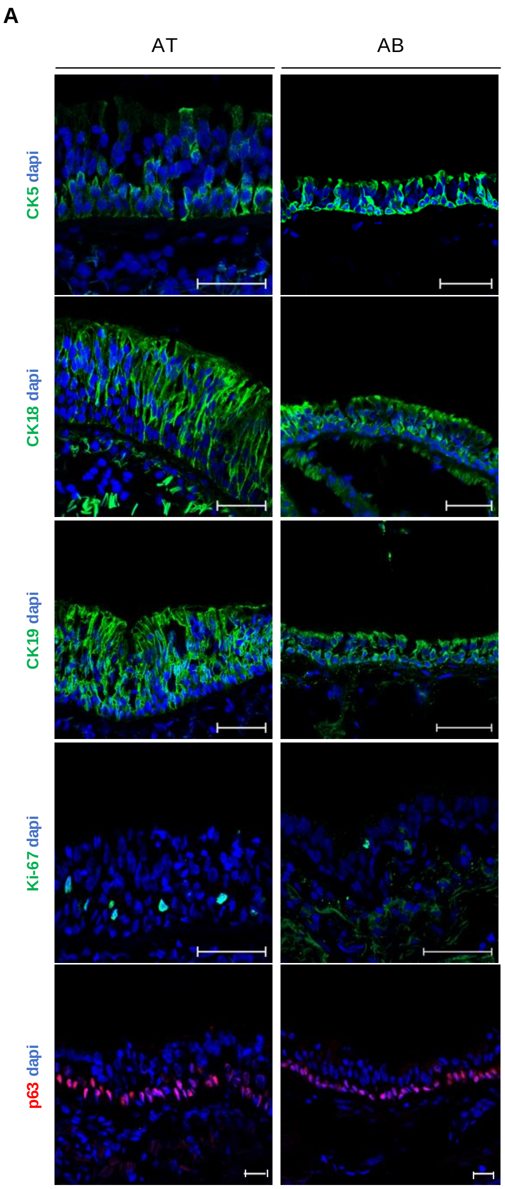

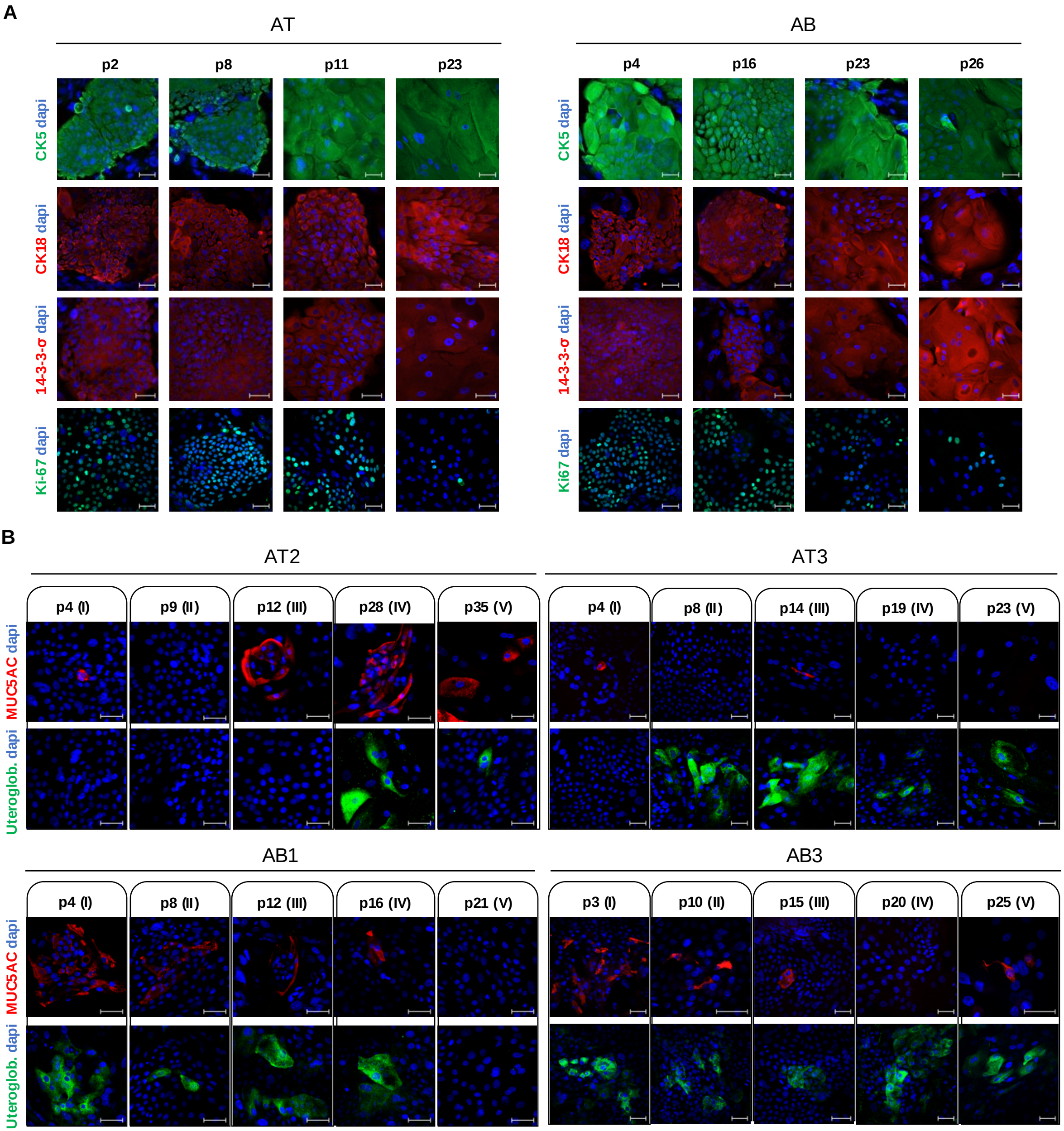

**A**

| | H | M | P |
|---|---|---|---|
| **AT clonal types (%)** | 8.7 ± 2.3 | 77.1 ± 3.7 | 14.2 ± 5.1 |
| **AB clonal types (%)** | 23.7 ± 14.4 | 71.7 ± 9.7 | 4.6 ± 4.6 |

#### Figure S4

**B**

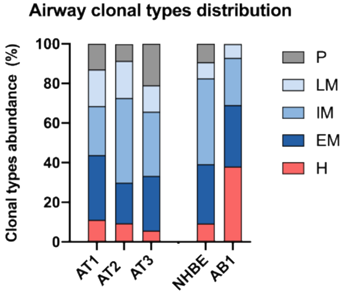


**C**

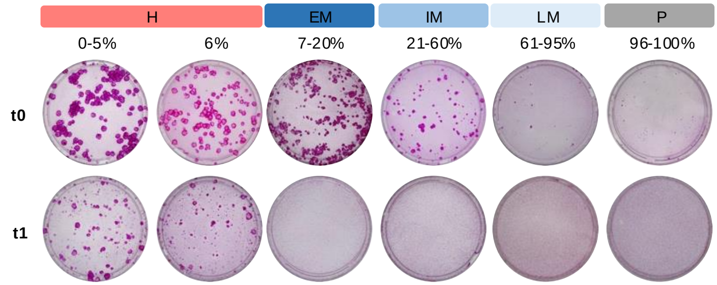


**D**

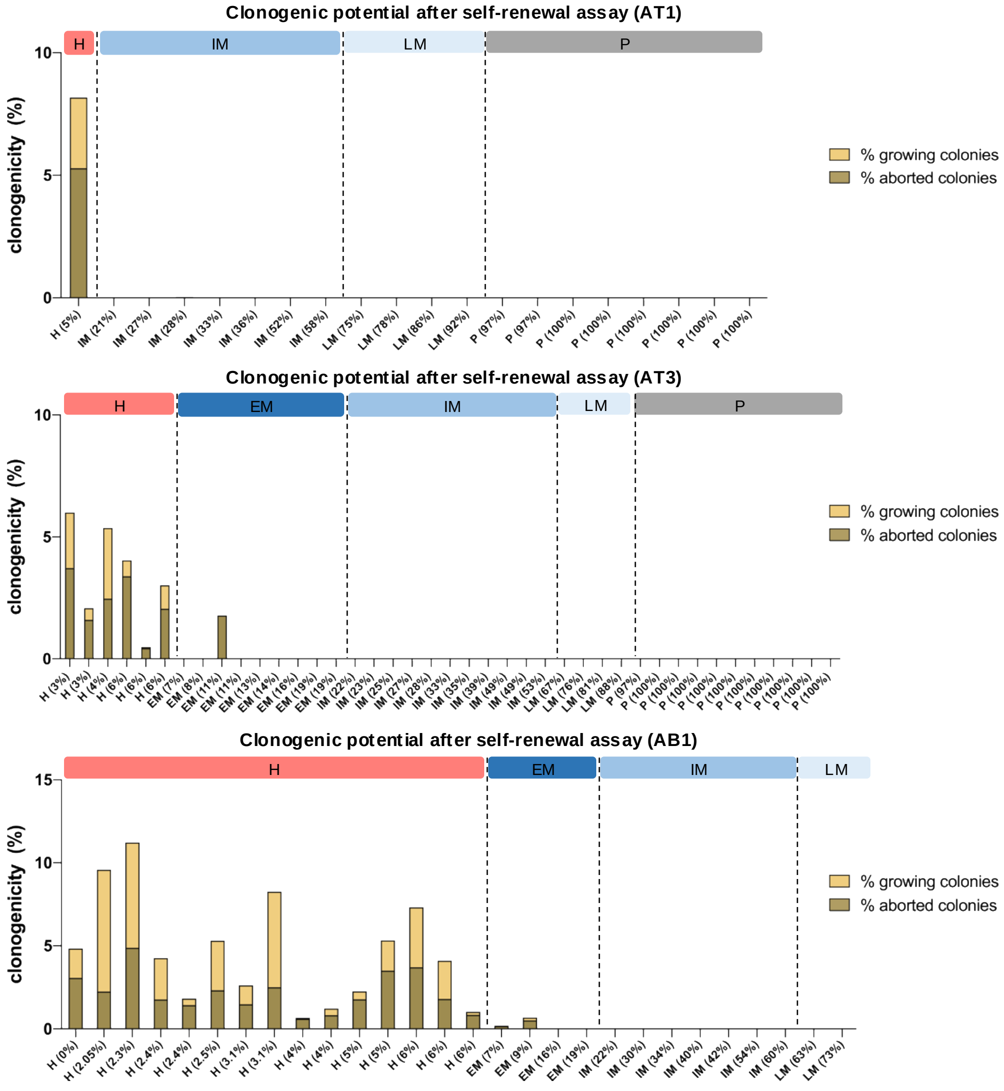

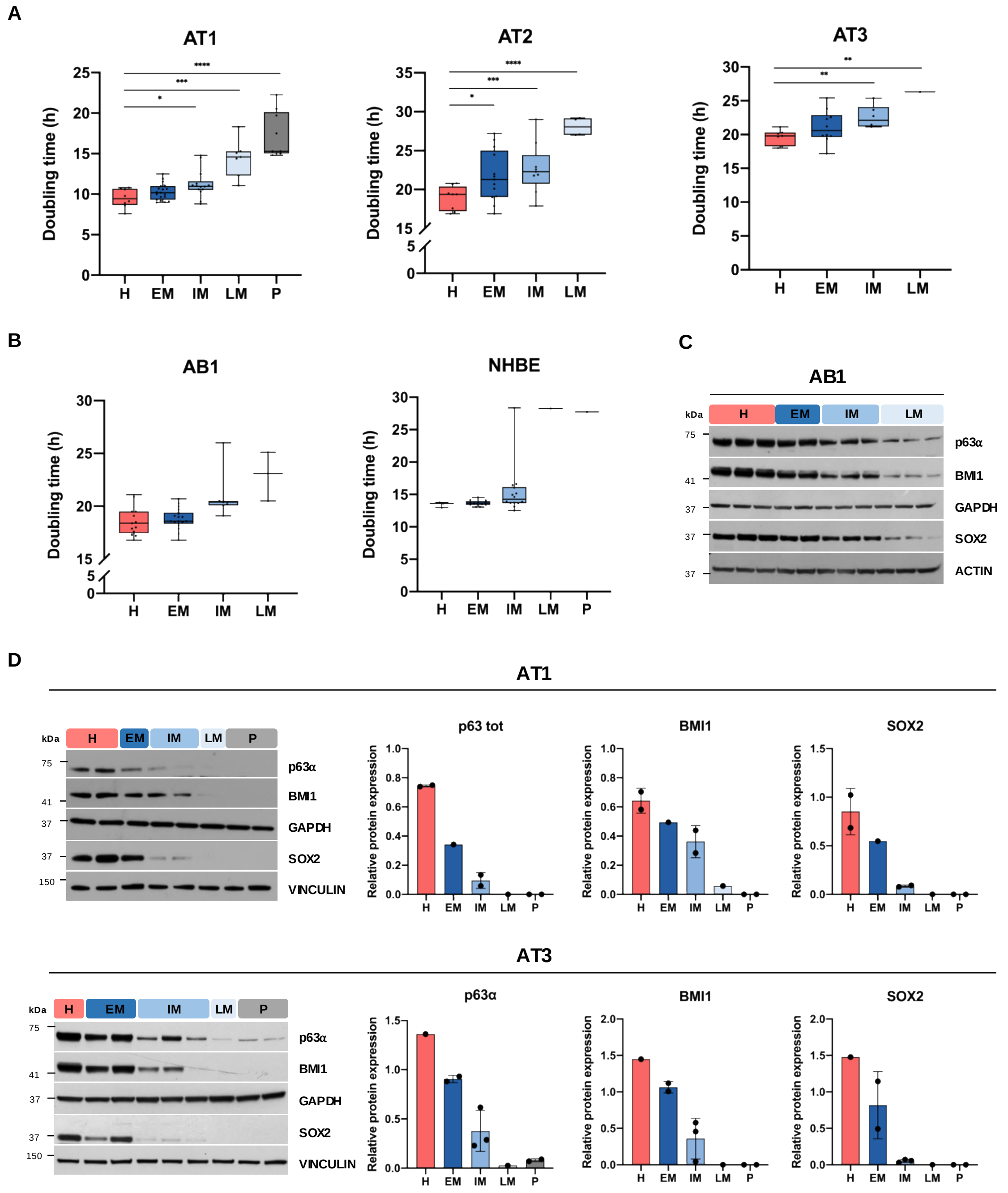
